# Gut microbiome stratification reveals functional metabolic signatures linked to host health

**DOI:** 10.64898/2026.05.05.722951

**Authors:** Mathilde Sola, Anne Hiol, Giacomo Vitali, Sébastien Fromentin, Marine Gilles, Emmanuelle Le Chatelier, Christian Morabito, Florian Plaza Oñate, Nicolas Pons, Benoît Quinquis, Florence Thirion, Jordan Denis, Renaud Léonard, Corinne Cruaud, Patrick Wincker, Pedro H. Oliveira, Guilhem Sommeria-Klein, Le French Gut Consortium, Catherine Robbe Masselot, Mathieu Almeida, Hervé Blottière, Joël Doré, Dusko S. Ehrlich, Robert Benamouzig, Clémence Frioux, Magali Berland, Patrick Veiga

## Abstract

The human gut microbiome exhibits reproducible taxonomic configurations, yet the ecological processes linking them to health remain unclear. We applied the five-enterosignature (ES) model to 5,169 metagenomes of *Le French Gut* participants and developed a rule-based framework that stratifies continuous microbiome variation into interpretable partitions. Across these partitions, hydrogen disposal pathways formed distinct functional gradients, reproduced in an international meta-cohort (n = 5,107). Partitions dominated by *Prevotella* enterosignature showed greater sulfate-reduction potential, *ES-Firmicutes*-associated partitions greater methanogenesis potential, and *ES-Bacteroides*-enriched partitions greater acetogenesis potential. The latter were also characterized by oxidative-stress and mucin-degradation signatures, altered fecal mucin O-glycosylation, and less favorable health indicators, including more frequent digestive symptoms, higher perceived stress, and more psychotropic drug prescriptions. These patterns were also accompanied by distinct dietary profiles. Our findings link hydrogen metabolism, redox-associated functions, mucin utilization, diet, and host health, and support enterosignature-based partitioning for cross-cohort comparisons and future stratified interventions.

## 1 Introduction

The gut microbiome is a key regulator of host physiology and is strongly linked to human health and disease, including cardiometabolic and inflammatory conditions, as well as mental disorders via the gut-brain axis [1–3]. Across ecosystems, biotic and abiotic forces jointly shape community structure and can lead to stable ecological configurations organized along functional gradients [4–6]. Yet, despite extensive study and the availability of large-scale metagenomic datasets, the ecological drivers of the gut microbiome structure remain unclear, with host, dietary, and lifestyle factors typically explaining only a limited fraction of inter-individual variability [7].

The identification of global structures in the gut microbiome – such as enterotypes, enterobranches, and enterosignatures – has demonstrated that the intestinal ecosystem is not randomly organized but follows a reproducible architecture [8–10]. These frameworks converge on three major ecological configurations in adults, characterized by the dominance of *Bacteroides*, *Firmicutes* (Bacillota) or *Prevotella* but differ in their scalability and conceptual assumptions. While enterotypes rely on discrete clustering [8] and enterobranches require very large cohorts [9], the enterosignature (ES) model represents gut microbial compositions as combinations of five generalizable ecological signatures, thereby better capturing gradual variation [10]. These five ES accurately described samples beyond the original training dataset, including samples from non-Western populations [10], and corresponding signatures were identified using distinct metagenomic approaches, supporting the robustness and potential reusability of the model [11]. These frameworks have proven useful to describe population-level organization; yet, two fundamental questions remain insufficiently addressed: which ecological and functional features characterize the resulting microbiome configurations, and how are these configurations associated with host health?

Understanding the relationships between gut microbiome structure and health can provide therapeutic leverage, but it requires large, well-characterized datasets capable of capturing inter-individual variability. One option is to aggregate small cohorts; however, this often introduces biases and limitations in data consistency and quality, which can affect the reliability and interpretability of observed associations [12]. The *Le French Gut* (LFG) project [13], beyond being a large cohort, exemplifies homogeneous and standardized acquisition of both biological data and extensive health, diet and lifestyle data, thereby minimizing methodological biases and opening the way for more robust analyses and interpretations.

In this study, we analyzed the first 5,169 participants in the LFG cohort from a microbial ecology perspective to investigate the functional organization underlying variation in gut microbiome composition and identify the associated environmental and host factors. Using the ES framework, we stratified the gut microbiome continuum into seven main partitions reflecting both dominant and subdominant ecological signals. Hydrogen metabolism emerged as a prominent ecological axis differentiating these strata, a pattern further validated in an independent international meta-cohort. We then integrated fecal mucin O-glycosylation profiles with extensive health, lifestyle, and dietary metadata to examine the relationships between this ecological organization and host physiology and well-being. Together, our findings highlight the value of enterosignature-based partitioning for investigating gut microbiome ecology and identify hydrogen metabolism as a potentially central dimension of microbial organization, possibly reflecting metabolic trade-offs with implications for host health.

## 2 Results

### 2.1 The *Le French Gut* cohort

The LFG project launched a nationwide cross-sectional study in France, aiming to collect up to 100,000 fecal samples from the general population. The study integrates detailed health, lifestyle, and dietary data via self-administered questionnaires, and is further enriched through linkage with national health databases (*Système National des Données de Santé*, SNDS) and environmental national resources, including regional water composition databases [13].

For the present study, we analyzed data from the first 5,169 participants for whom shotgun metagenomic profiling was available. These participants included 66 % women, covered a wide range of ages (from 18 to 87 years old) and body mass indices (BMI), and included 28 % of participants diagnosed with at least one chronic disease. The main characteristics of the cohort are summarized in the Extended Table 1. Overall, this cohort captures heterogeneous demographic and health profiles and provides an opportunity to explore the ecological and functional features associated with gut microbiome configurations.

### 2.2 Partitioning the *Le French Gut* cohort to delineate an ecological continuum

To further characterize the ecological landscape of our cohort, we applied the genus-level five-enterosignature (ES) model developed by Frioux *et al.* [10] to the abundance profiles of the LFG cohort. Each sample was described by the relative contributions of five ES: ES-*Bifidobacterium*, ES-*Escherichia*, ES-*Bacteroides*, ES-*Firmicutes* (Bacillota), and ES-*Prevotella* (Fig. 1a). Examination of the primary ES (*i.e.*, the ES with the largest relative contribution) showed that the latter three are the predominant primary signatures in the cohort. ES-*Firmicutes* was the primary ES in 60 % of samples, consistent with previous general-population studies [8, 10]. Although each sample was represented as a combination of five ES, two or three ES were sufficient to reach a cumulative contribution of 90 % in most samples (see Methods), consistent with previously reported patterns [10]. In contrast, this threshold was reached by a single ES in only 3.5 % of samples, most often ES-*Prevotella* (Fig. 1b), supporting the relevance of mixture models for representing gut microbiome composition.

**Extended Table 1.** Characteristics of *Le French Gut* participants. Abbreviations: BMI, Body Mass Index; DSFQ, Digestive Symptom Frequency Questionnaire.

| **Characteristics** |  | **Mean [min - max] (%)** |
|---|---|---|
| Participants | Total number | 5,169 |
|  | Optional food survey | 1,696 (32.7 %) |
| Age (years old) |  | 49.0 [18 - 87] |
| Sex | Female | 3,431 (66.4 %) |
|  | Male | 1,738 (33.6 %) |
| BMI (kg/m<sup>2</sup>) |  | 24.3 [12.9 - 73.9] |
| Chronic disease diagnosed | Yes | 1,453 (28.1 %) |
|  | No | 3,568 (69.0 %) |
|  | ”I don’t know” | 148 (2.9 %) |
| Smoking habit | Smokers | 586 (11.3 %) |
|  | Former smokers | 1,181 (22.8 %) |
|  | Non smokers | 3,402 (65.8 %) |
| Digestive Symptom Frequency (DSFQ) [14] |  | 5.75 [0 - 16] |
| Bristol Stool Scale score |  | 3.83 [1 - 7] |
| Perceived Stress score (PSS-4) [15] |  | 4.83 [0 - 16] |

**Fig. 1:**
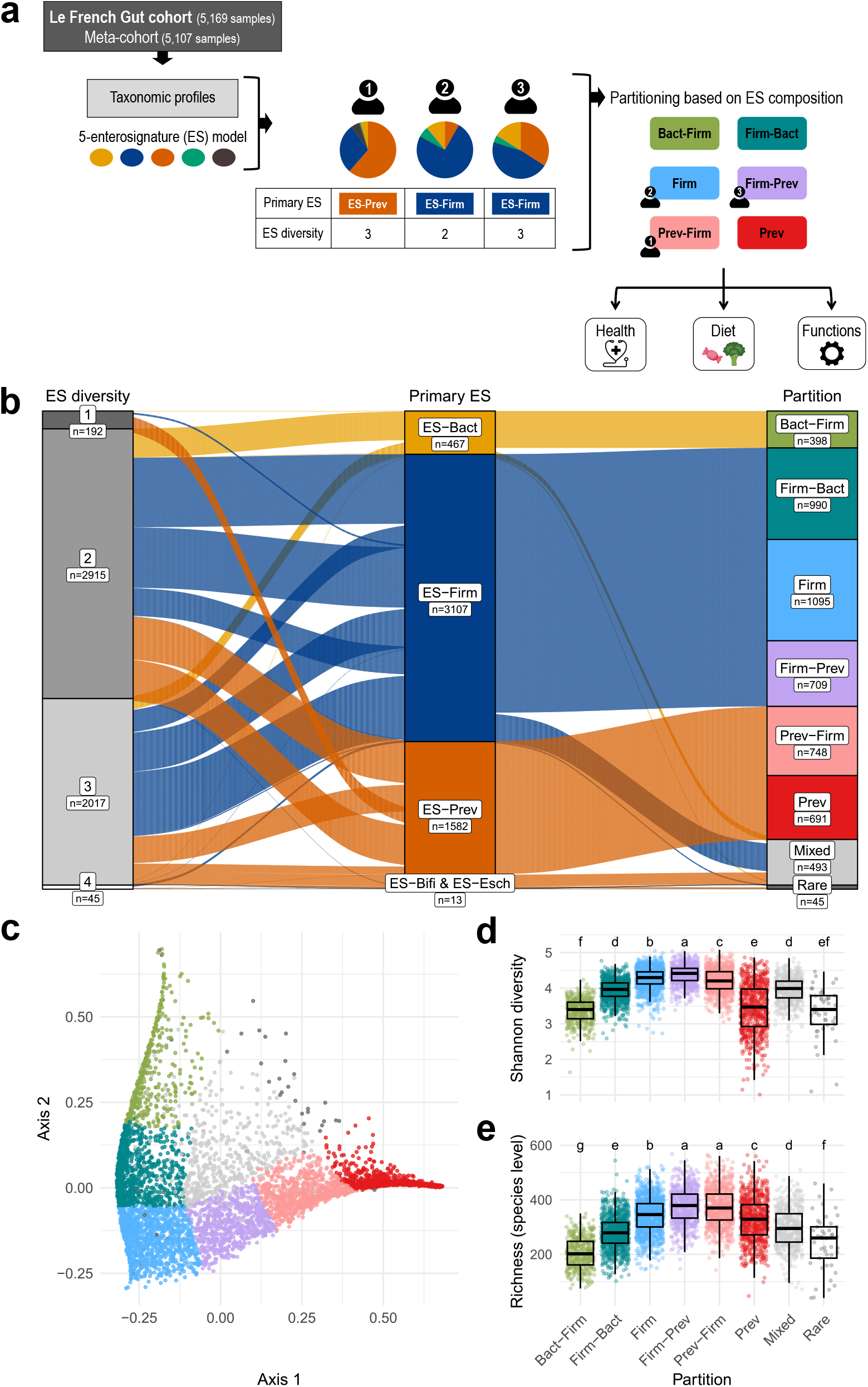
Enterosignature-based stratification of the LFG cohort. a) Description of the analysis workflow. b) Individuals distribution across ES and their corresponding partitions. ES diversity refers to the number of prominent ES in each sample (see Methods) c) Principal Coordinates Analysis of Bray-Curtis dissimilarities based on the ES composition of the LFG samples. Each sample is colored according to its partition. d) Distribution of the Shannon diversity of the LFG samples. e) Distribution of the richness of the LFG samples calculated at the species level. Letters correspond to the result of Tukey’s HSD post-hoc test to assess pairwise significant differences. Different letters indicate that groups are significant^3^l^9^y different.

To group samples with similar ES compositions, we developed a generalizable rule-based stratification procedure based on the relative dominance of the primary ES and the combined contribution of the primary and secondary ES (see Methods, Fig. 1a,b, and Supp. Figs. S1 and S2). Briefly, samples with a primary-to-secondary ES ratio of at least 3 were considered relatively dominated by their primary ES. Among the remaining samples, those in which the primary and secondary ES jointly contributed at least 80% were assigned to an ordered two-ES partition, whereas all other samples were assigned to the *Mixed* partition.

This procedure discretized the gradual transitions from ES-*Bacteroides* to ES-*Prevotella* through ES-*Firmicutes* into the following partitions: *Bact-Firm*, *Firm-Bact*, *Firm*, *Firm-Prev*, *Prev-Firm*, and *Prev* (Fig. 1c). Partition names indicate the primary ES and, for two-ES profiles, the secondary ES. For example, the *Prev-Firm* partition includes samples in which ES-*Prevotella* is primary and ES-*Firmicutes* makes a substantial secondary contribution, whereas the *Prev* partition includes samples in which ES-*Prevotella* is strongly dominant relative to the secondary ES. The procedure also identified a *Bact* partition comprising 73 individuals (1.4% of the cohort), which was merged with *Bact-Firm* for downstream statistical analyses because of its small size. Other rare ES profiles (1 *≤ n ≤* 16) were pooled into a separate *Rare* category. After merging *Bact* with *Bact-Firm*, the six compositionally defined partitions and the *Mixed* partition constituted the seven main partitions considered in the LFG cohort.

Applying the same procedure to the dataset used to define the five-ES model of Frioux *et al.* [10] identified three additional partitions, reflecting the broader diversity of microbiome profiles represented in that population (Supp. Fig. S2d).

The distributions of ES contributions across partitions in the LFG cohort are shown in Supp. Fig. S2b. The *Mixed* partition comprised 9.5% of samples and displayed relatively balanced contributions from several ES. Because this partition was compositionally heterogeneous (Supp. Fig. S1c), it was excluded from subsequent analyses.

The *Bact-Firm*, *Firm*, and *Prev* partitions were selected as three archetypal ES configurations spanning the compositional continuum: *Bact-Firm* and *Prev* occupied opposite ends of the gradient visualized by PCoA, whereas *Firm* occupied a central position. Differential abundance analysis identified the main taxa distinguishing these partitions. Characteristic metagenomic species (MSPs) included *Flavonifractor plautii*, *Phocaeicola vulgatus*, and *Bacteroides uniformis* in *Bact-Firm*; *Maccoya intestinihominis*, *Scatomorpha sp900545405*, and *Lawsonibacter A hominis* in *Firm*; and *Prevotella copri B*, *Suilimivivens aceti*, and *Suilimivivens sp000438155* in *Prev* (Supp. Fig. S3 and Supp. Table 1). Their abundances shifted gradually across partitions, supporting the interpretation that these partitions discretize an underlying microbial continuum rather than represent discrete ecological states.

Alpha-diversity varied significantly across partitions along the ES continuum (Fig. 1d,e). *Firm*, *Firm-Prev*, and *Prev-Firm* partitions exhibited the highest diversity, with mean species richness reaching 376 in *Firm-Prev*, whereas *Bact-Firm* showed lower values with a mean richness of 205. Shannon diversity also differed significantly across all partitions along the same trend (Kruskal-Wallis q-value *<* 2.2e-16). Notably, Shannon diversity within the *Prev* partition spanned a wide range of values (Fig. 1d), suggesting internal heterogeneity or subpopulations. Stratifying *Prev* individuals by median Shannon index showed that low-diversity profiles were strongly dominated by *Prevotella copri B*, whereas high-diversity profiles exhibited lower dominance of *P. copri B* and enrichment in *Oscillospiraceae* and *Christensenellales* (Supp. Fig. S4). A strain-level analysis further showed that low- and high-diversity groups harbored distinct *P. copri B* strains, indicating that diversity differences within the *Prev* partition are associated with strain-level variation rather than taxonomic composition alone (Supp. Fig. S4).

### 2.3 Dihydrogen recycling routes and functional potential across partitions

We next examined the functional modules that most strongly distinguished the archetypal partitions *Bact-Firm*, *Firm* and *Prev*, selecting those showing at least a 10 % difference in prevalence between any two partitions (Methods, Supp. Table 2). Modules associated with the major gut H_2_-consuming routes (main pathways illustrated in Supp. Fig. S5) — methanogenesis, dissimilatory sulfate reduction, and acetogenesis — were among the strongest discriminatory signals. Their prevalence peaked towards the *Firm*, *Prev*, and *Bact-Firm* partitions, respectively (Fig. 2a). Because sex differences in methanogenesis have previously been reported [16], we performed sex-stratified analyses and observed similar trends in male and female participants (Supp. Fig. S6), thereby allowing the pooling of both sexes. The hydrogenotrophic potential of the sulfate reduction and acetogenesis-associated modules was further supported by the identification of H_2_-utilization genes in MSPs carrying the corresponding modules, including *Desulfovibrio* MSPs and *Blautia A caecimuris* (Supp. Fig. S7).

**Fig. 2:**
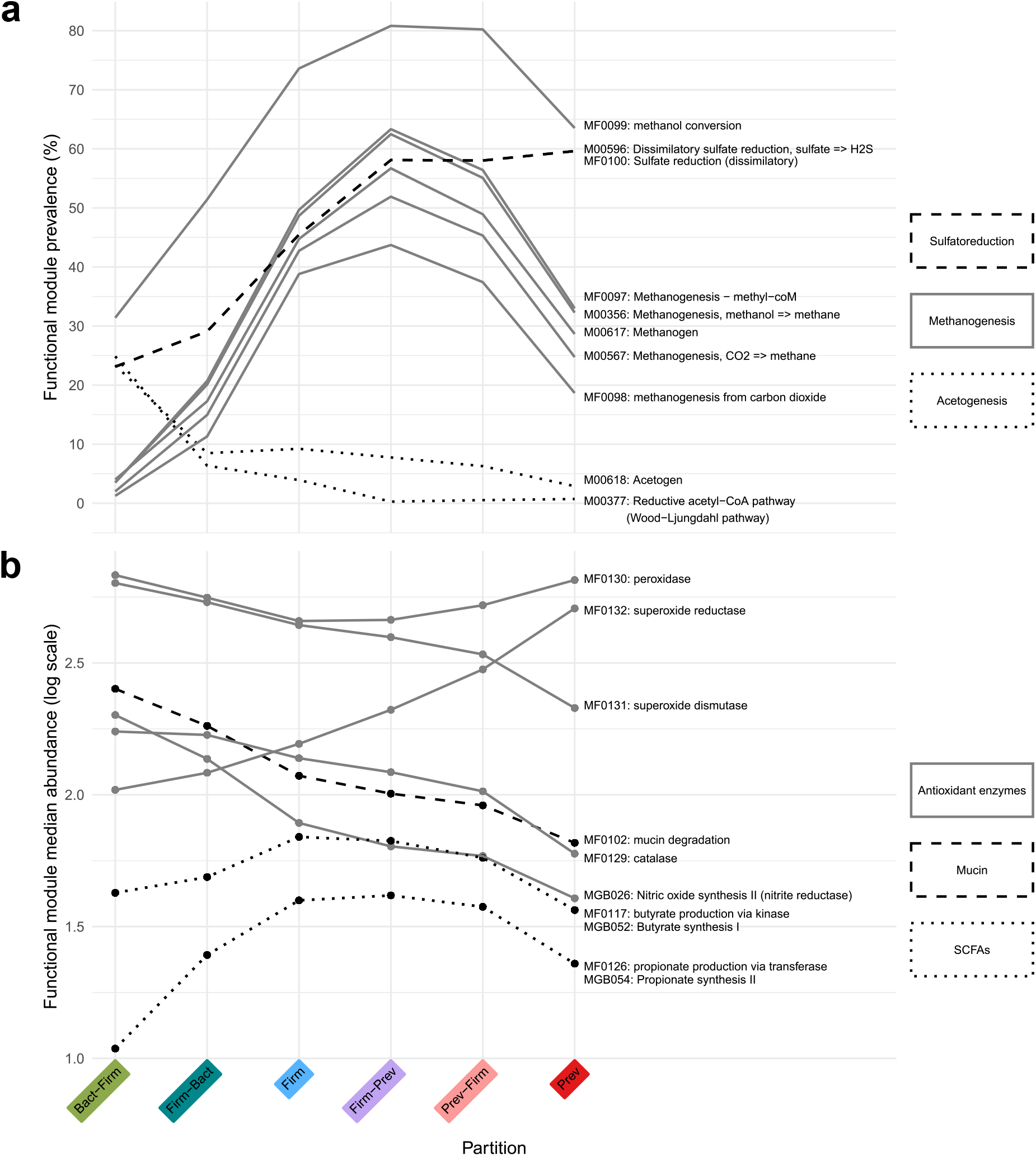
Functional analysis reveals distinct metabolic patterns across partitions. a) Prevalence of functional modules linked to dihydrogen metabolism across partitions. These modules belong to three categories: sulfatoreduction, acetogenesis, and methanogenesis. b) Abundance of functional modules across partitions. These modules are linked to antioxidant enzymes, mucin degradation and production of butyrate and propionate. The median is plotted. Abreviations: SCFAs, Short Chain Fatty Acids

To complete our functional exploration, we focused on differentially abundant functional modules across microbiome partitions. Among the 122 statistically significant modules identified (Kruskal-Wallis test q-value *<* 0.05 and Cliff’s Delta (*δ*) *>* 0.6, full list in Supp. Table 3), reactive oxygen species (ROS) related functions displayed a clear gradient (Fig. 2b). We observed a continuous shift from superoxide dismutase-dominated (SOD, maximum Cliff’s Delta *δ_max_* = 1.0) to superoxide reductase-dominated (SOR, *δ_max_* = -0.99) configurations, progressing from the *Bact-Firm* to the *Prev* partitions, accompanied by a decrease in catalase (*δ_max_*= 0.86) and nitrate reductase (*δ_max_*= 0.94) module abundances. SOD and catalase detoxify ROS while producing molecular oxygen, whereas SOR and peroxidases detoxify ROS without oxygen production, suggesting a shift from oxygen-dependent to oxygen-independent redox strategies. The *Bact-Firm* partition showed increased abundance of the mucin-degrading module, whereas butyrate and propionate biosynthesis pathways decreased from the *Firm* partition to the *Bact-Firm* and *Prev* partitions, indicating differences in predicted fermentative potential (Fig. 2b).

These patterns were also observed in regression analyses relating module abundances to continuous enterosignature proportions, supporting their robustness beyond partition-based comparisons (Supp. Fig. S8). When visualizing the contribution of each ES to the prediction of module abundance, we observed for instance, that the predicted abundance of the mucin degradation module (MF0102) increases as the proportion of *ES-Bact* increases.

Together, our results reveal gradual functional shifts along the partitioning gradient, including hydrogen disposal pathways, reactive oxygen species-related processes, and mucin degradation.

### 2.4 Exploration of fecal mucin O-glycan profiles support partition-specific mucus alterations

To determine whether the mucin-related functional signal identified by metagenomic profiling was associated with measurable biochemical differences in fecal mucus, we analyzed mucin O-glycans in 26 stool samples distributed across 3 partitions (*Bact-Firm*, *n* = 7; *Firm*, *n* = 7; and *Prev*, *n* = 12), as fecal mucin glycosylation has been shown to reflect proximal colonic mucus profiles [17, 18]. Human intestinal mucins are mostly composed of acidic O-glycans, most of which are built on a core 3 scaffold (GlcNAc*β*1-3GalNAc) and frequently carry an *α*2,6-linked sialic acid on the initial GalNAc [18] (Fig. 3a).

**Fig. 3:**
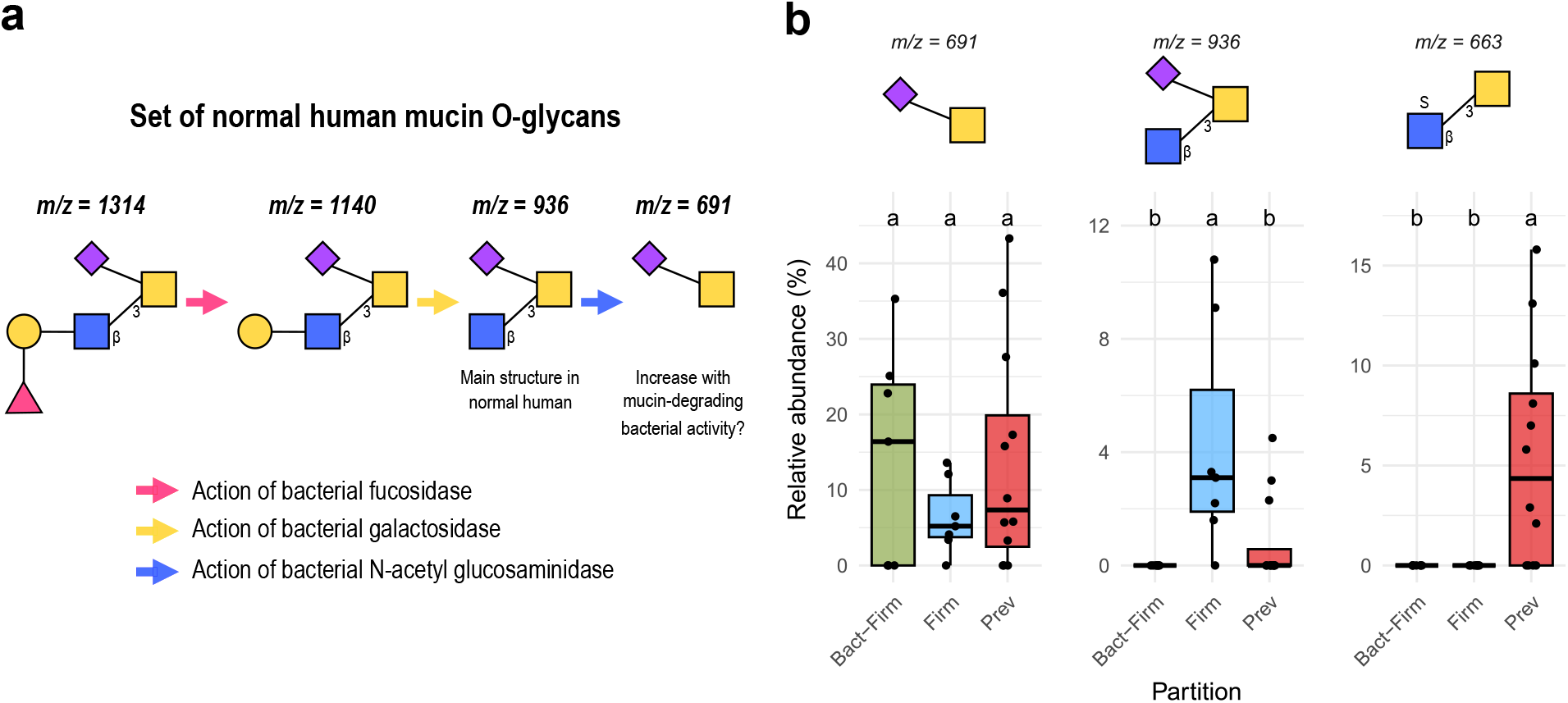
Exploratory fecal mucin glycomics reveals partition-specific patterns of sialylation and sulfation across partitions. a) Normal human mucin glycan expression patterns and degradation products of mucin-degrading bacteria. b) Relative abundances (%) of selected mucin O-glycan ions. Letters correspond to the result of Tukey’s HSD post-hoc test to assess pairwise significant differences. Different letters indicate that groups are significantly different.

Against this structural background, fecal mucin glycomics revealed clear differences in the distribution of sialylated glycans across partitions (Fig. 3b, Supp. Table 4). The ion at m/z 691, corresponding to a NeuAc residue *α*2,6-linked to the GalNAc at the attachment site, was more abundant in *Bact-Firm*. This pattern is consistent with degradation of the 3-linked branch while preserving a residual *α*2,6-sialylated structure on the core GalNAc. In contrast, *Firm* samples retained higher levels of the ion at m/z 936, corresponding to a sialylated core 3 structure, which is the major glycan expressed in human intestinal mucins. *Prev* samples included low and high diversity members reflecting the subgroups mentioned in section 2.2. Low-diversity *Prev* mucin profiles also displayed detectable m/z 936, together with lower m/z 691 than *Bact-Firm*, indicating a distinct sialylation pattern, whereas the profile in high-diversity *Prev* samples indicated more degradation of sialylated structures. Sulfated glycans also differed across partitions: the ion at m/z 663, corresponding to a sulfated GlcNAc linked in position 3 to the initial GalNAc, was enriched in *Prev*, indicating a greater representation of sulfated mucin motifs.

Together, these data provide biochemical support for the partition-specific mucus signal identified by metagenomics, linking the *Bact-Firm* partition to stronger mucin glycan remodeling and the *Prev* partition to a more sulfated mucin landscape.

### 2.5 Replication on an international meta-cohort

To assess the generalizability of partitions on taxonomic and functional compositions, we applied the same computational analyses to an international meta-cohort (5,107 samples). This meta-cohort comprises eight curated cohorts from nine industrialized countries (see Methods), with metadata distributions comparable to the LFG cohort (Fig. 4a, Supp. Fig. S9). Mean age was 51.7 years and mean BMI 26.6 kg/m^2^; women represented 60.8 % of participants, and 30.7 % reported a chronic disease.

**Fig. 4:**
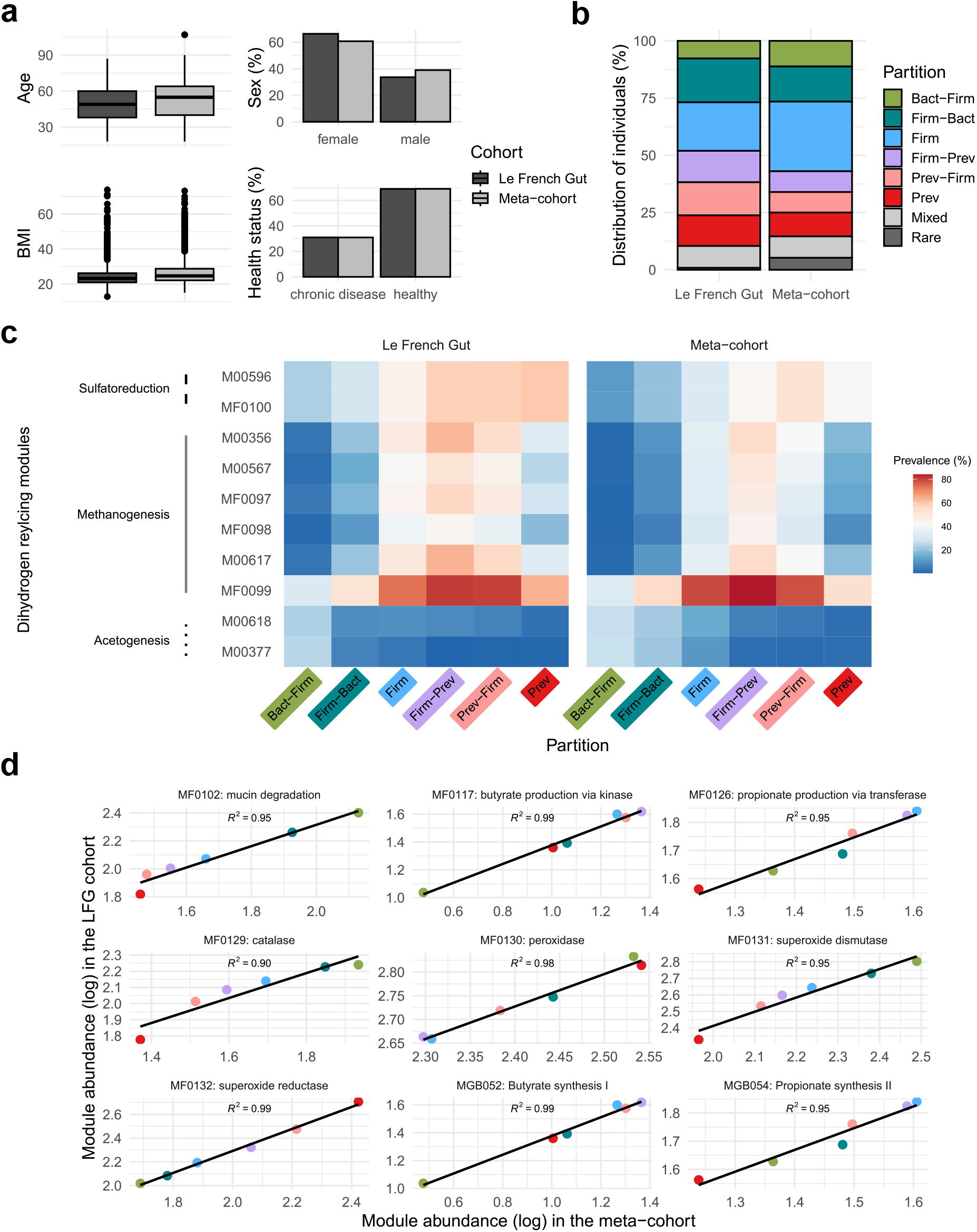
Replication in the international meta-cohort confirms patterns observed in the LFG cohort. a) Mirroring of minimal metadata between the two cohorts. b) Distribution of individuals across partitions in both cohorts. c) Heatmaps of the prevalence of functional modules related to H_2_ recycling across partitions for both cohorts. d) Comparative abundance plot of the functional modules that emerged in Fig. 2b between the two cohorts. Median abundance for each partition is plotted.

ES-based partitioning yielded a similar distribution of individuals (Fig. 4b, Supp. Fig. S10), with a higher proportion in the *Firm* partition, indicating that ES-based stratification is consistent across industrialized populations regardless of country. The *Bact* partition was more prevalent in the metacohort (*n* = 183) and exceeded the 2% threshold used to define rare partitions (Methods). Nevertheless, it was merged with the *Bact-Firm* partition to maintain a consistent partitioning scheme and enable direct comparisons with the LFG cohort.

Functional module prevalence analysis across partitions in the meta-cohort highlighted again H_2_-related pathways. Comparison between both datasets showed nearly identical prevalence profiles for sulfatoreduction, methanogenesis and acetogenesis modules (Fig. 4c, Supp. Fig. S11, Supp. Table 5). Differentially abundant modules identified above were also significant in the meta-cohort (Fig. 4d, Supp. Fig. S11, Supp. Table 6). Median abundances per partition were strongly correlated between datasets, with linear regression R^2^ values close to 1. Overall, replication of partitioning and functional analyses confirms the robustness and generalizability of our findings across distinct populations.

### 2.6 Diet and lifestyle vary across gut microbiome partitions

We next examined the dietary context of the partitions to identify dietary features potentially relevant to the functional modules that distinguished them. We first focused on dietary fiber, whose microbial fermentation can generate H_2_ and whose limited availability may favor microbial mucin degradation [19], using data from the mandatory (short) dietary questionnaire (*n* = 5, 169, Fig. 5a). Mean fiber intake was lower in *Bact-Firm* individuals than in the *Prev-Firm* partition (Mann-Whitney U test p-value 4.19e-5). Similarly, annual Fruit and Vegetables Diversity (the number of different items consumed over a year) was higher in *Firm-Prev* and lower in *Bact-Firm*.

**Fig. 5:**
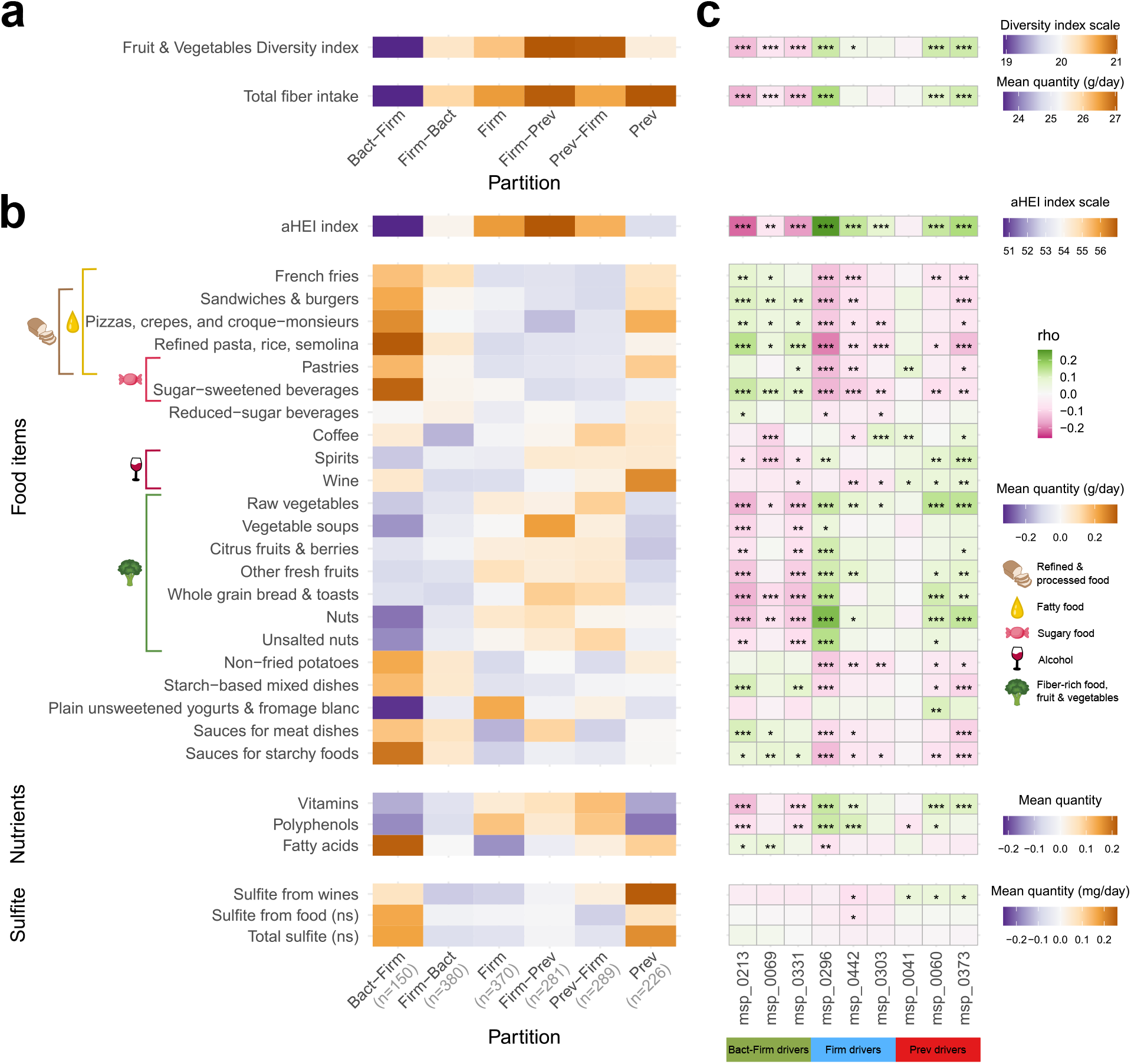
Dietary intake is significantly distinct between partitions. a) Focus on fiber intake from the general questionnaire answered by all LFG participants (*n* = 5, 169) across partitions. b) Nutritional profiles across partitions of LFG participants who completed the FFQ questionnaire (*n* = 1, 696) covering global index of dietary quality, intake of food items, nutrients, and sulfite. Nutritional variables were centered and scaled. All variables showed significant differences across partitions (Kruskal-Wallis test q-value *<* 0.05), except those labeled “ns”. The color hues correspond to the mean value of each sub-heatmap. c) Correlations between partitions species drivers and dietary patterns. The top 3 drivers per partition were identified across the three archetypal partitions (see Methods). The ranking was obtained by ordering their mean cliff’s delta in absolute value. Spearman correlations were corrected for FDR; the tile color corresponds to the Spearman’s rho value; the significance level is labeled with stars: * *<*0.05, ** *<*0.01, *** *<*0.001. Identified drivers: msp_0213: *Flavonifractor plautii*, msp_0069: *Phocaeicola vulgatus*, msp_0331: *Oscillobacter welbionis*, msp_0296: *Maccoya intestinihominis*, msp_0442: *Lawsonibacter A hominis*, msp_0303: *Scatomorpha sp900545405*, msp_0041: *Prevotella copri B*, msp_0060: *Suilimivivens sp000438155*, msp_0373: *Suilimivivens aceti*.

We then investigated differences in the alternative Healthy Eating Index (aHEI) across partitions, given its reported strong association with gut microbiome composition [20]. For this analysis, we used Food Frequency Questionnaire (FFQ) data from the subset of participants who voluntarily completed the questionnaire (*n* = 1, 696, Fig. 5b). Higher diet quality was associated with the *Firm*, *Firm-Prev*, and *Prev-Firm* partitions, with progressively lower scores towards the *Bact-Firm* and *Prev* partitions.

Participants in *Firm*-dominant partitions reported greater consumption of foods generally considered beneficial, including raw vegetables, vegetable soups, fresh fruits, nuts and whole-grain bread and toasts, reflected in higher polyphenol and vitamin intake. In contrast, the *Bact-Firm* partition – and to a lesser extent the *Prev* one – were characterized by higher consumption of energy-dense foods, including French fries, sandwiches and burgers, pizzas, and crepes, alongside lower vegetable consumption, reflected in higher fatty acid intake. *Bact-Firm* individuals additionally showed higher consumption of pastries, sugar-sweetened beverages, and sauces, together with lower consumption of yogurts, whereas the *Prev* partition reported greater wine intake. All of these variables differed significantly across partitions (q-values *<* 0.05, Supp. Table 7).

Prompted by the enrichment of sulfate-reduction pathways in *Prev*, we investigated potential dietary and waterborne sources of sulfate and sulfite that could support sulfate reduction. Individual sulfite exposure derived from FFQ data was markedly higher in the *Prev* partition, largely driven by wine consumption (Fig. 5b). Sulfate intake from tap water, estimated by linking residential codes to the national water composition registry, was modestly higher in *Prev* -dominated partitions, but differences were not statistically significant (Supp. Fig. S12).

To assess whether partition-level dietary patterns were associated with species-level taxonomic differences, we analyzed correlations between dietary variables and the abundances of taxonomic drivers, defined as the MSPs showing the largest abundance differences among the archetypal partitions (*Bact-Firm*, *Firm* and *Prev*), and dietary variables (Supp. Table 8). We focused on the three main taxonomic drivers per partition (Fig. 5c). Eight of the nine drivers showed associations with aHEI consistent with the dietary profile of their respective partition. Interestingly, all three *Firm* partition drivers were associated with aHEI, but only *Maccoya intestinihominis* correlated with fiber intake, likely reflecting its association with the consumption of plant-based foods captured by this index. The other two, *Lawsonibacter A hominis* and *Scatomorpha sp900545405*, may instead be associated with aHEI through other dimensions of dietary quality, such as sugar-sweetened beverages.

Associations with individual food items were heterogeneous: some drivers (*Flavonifractor plautii, Oscillobacter welbionis, Maccoya intestinihominis, Suilimivivens aceti*) correlated with more than 75 % of the 22 food items examined, whereas others (*Prevotella copri B* and *Scatomorpha sp900545405*) showed few associations.For MSPs associated with multiple food items, the directions of the correlations were often internally consistent: foods generally considered beneficial (*e.g.*, vegetables, fruits, and whole grains) tended to correlate in one direction, whereas less favorable dietary choices (*e.g.*, sandwiches and pizza) correlated in the opposite direction. Thus, these associations may reflect broader dietary patterns and the combined influence of multiple dietary components, rather than direct effects of individual foods or nutrients.

To extend lifestyle characterization beyond diet, we assessed weekly physical activity, which differed significantly across partitions (Chi2 test q-value 6.80e-5). Although overall patterns were similar, the *Firm-Bact*, and more markedly the *Bact-Firm* partitions, displayed a more sedentary profile (Supp. Fig. S13a). Finally, we constructed a composite Healthy Lifestyle Index (HLI) incorporating diet quality, physical activity, tobacco use, and alcohol consumption. A clear gradient emerged (Chi2 test q-value 6.35e-19): the *Bact-Firm* partition – and to a lesser extent the *Prev* one – had the lowest HLI scores, whereas the *Firm*, *Firm-Prev*, and *Prev-Firm* partitions showed progressively more favorable profiles (Supp. Fig. S13a).

### 2.7 *Bact-Firm* partition exhibits marked health alterations

To investigate whether partition-associated functions could help explain links with host health, we investigated associations between gut microbiome partitions, host- and health-related variables in the LFG cohort. Using self-reported survey data, we identified 186 variables significantly associated with ES-based partitions (q-values *<* 0.05, Supp. Table 9, Fig. 6a). Partitions differed in age, sex and BMI distributions (Fig. 6), with higher proportions of overweight and obese individuals in *Bact-Firm* and *Prev* partitions (Chi2 q-values of 1.72e-7 and 8.51e-7, respectively) and of participants older than 40 years old in the *Firm-Prev* group (Chi2 q-value 3.72e-3). In adjusted analyses, the reported number of chronic conditions was higher in *Bact-Firm* and *Firm-Bact* partitions than in the *Firm* one (adjusted incidence rate ratios with 95 % confidence intervals of 1.45 [1.17–1.80] and 1.26 [1.04–1.54], respectively). Perceived stress scores were also higher in *Bact-Firm* individuals (1.09 [1.01–1.18]), whereas differences in digestive symptom frequency (DSFQ) and sleep satisfaction were less pronounced, with confidence intervals overlapping 1. Distributions of these scores across partitions are illustrated in Supp. Fig. 13b. Associations with the *Bact-Firm* partition remained largely unchanged after additional adjustment for Bristol Stool Scale score, which varied significantly across partitions with the *Firm* individuals reporting lower values associated to constipation (Chi2 q-value 9.53e-5, Supp. Figs. S13b and S14). Associations with stress and digestive symptoms lost significance following additional adjustment for annual Fruit and Vegetables diversity, despite retaining similar trends, suggesting a potential contribution of diet to these inter-connected diet–microbiome–health relationships. Additional adjustment for weekly physical activity in the models had very little to no impact on the results (Supp. Fig. S14).

**Fig. 6:**
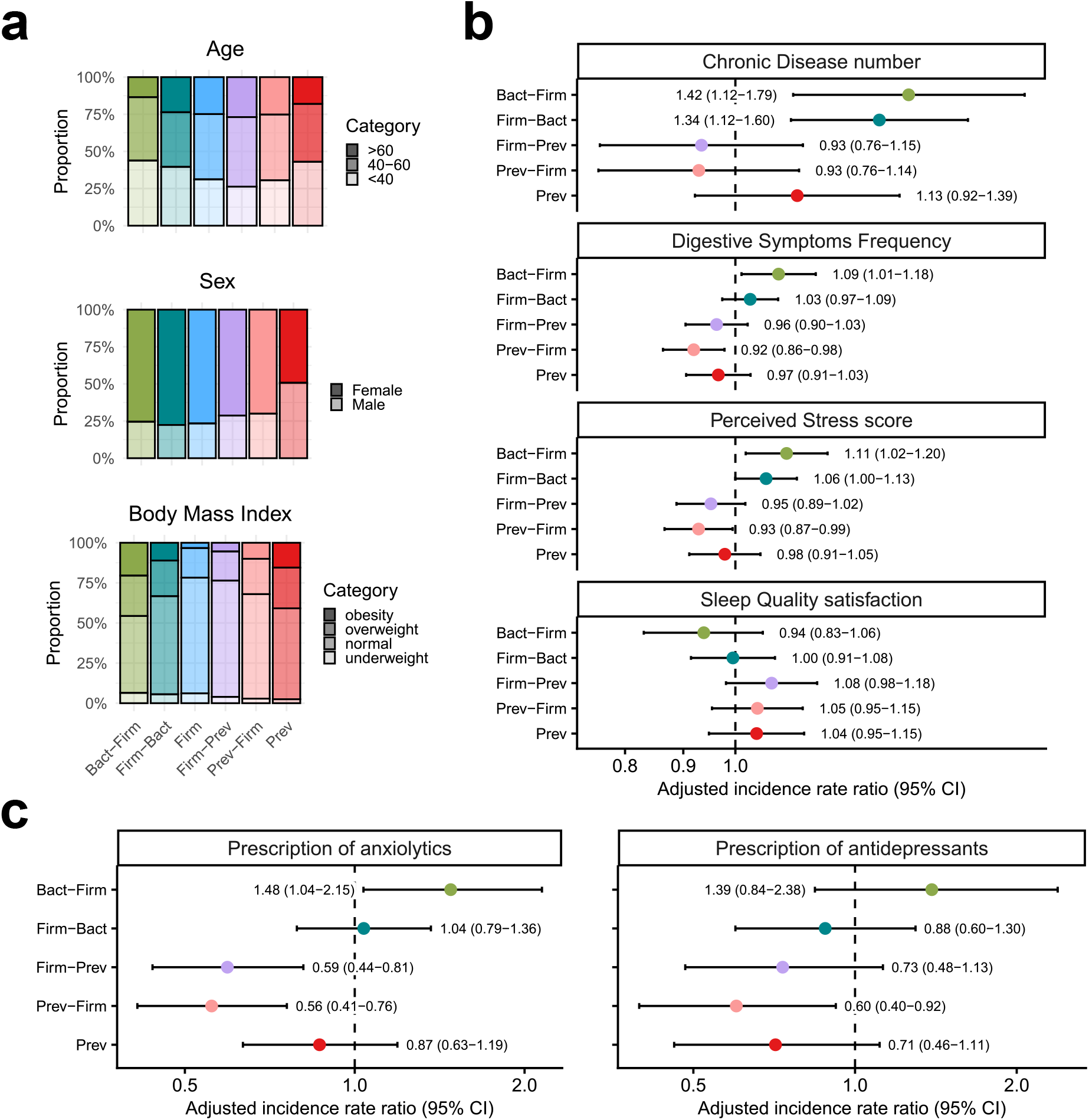
The health status of *Bact-Firm* and *Firm-Bact* partitions is altered. a) Distribution of participants across partitions for age, sex, and BMI. b) Associations between partitions and health-related outcomes, adjusted for age, sex, and BMI. Forest plots show adjusted incidence rate ratios (IRRs) and 95 % confidence intervals; the reference partition was *Firm*. The dashed vertical line indicates an IRR of 1. c) Anxiolytic prescriptions across partitions during the two years preceding participation. d) Antidepressants prescriptions across partitions during the two years preceding participation. Colors correspond to the partitions color code, transparency indicates the category level.

Prompted by differences in perceived stress, we leveraged linked data from the SNDS to examine prescriptions for anxiolytics and antidepressants across partitions in the two years preceding participation. Compared with the reference *Firm* partition, anxiolytic prescriptions were significantly more frequent among individuals in the *Bact-Firm* partition (1.48 [1.04–2.15]) and less frequent in the *Firm-Prev* and *Prev-Firm* partitions (0.59 [0.44–0.81] and 0.56 [0.41–0.76], respectively; Fig. 6c). A similar pattern was observed for antidepressant prescriptions, although the association was statistically significant only for the *Prev-Firm* partition (0.60 [0.40–0.92]). Consistent with these associations, the absolute prescription frequencies were highest in the *Bact-Firm* partition: 34% of participants had at least one anxiolytic prescription and 31% had at least one antidepressant prescription, a near twofold increase compared to the *Firm-Prev* and *Prev-Firm* partitions (16 % and 14 % for anxiolytics and antidepressants respectively), while *Firm-Bact* and *Firm* displayed intermediate levels (Supp. Fig. S13c). Similar patterns were observed when the prescription window was extended to 5 and 10 years before participation, although prescription records do not necessarily reflect actual drug use (Supp. Fig. S15).

Motivated by the marked mental health profile of the *Bact-Firm* partition, we revisited the 122 previously identified discriminant functional modules (Kruskal-Wallis q-value *<* 0.05; Cliff’s Delta *>* 0.6; Supp. Table 3) and focused on those annotated as gut-brain modules [21]. Several neuroactive metabolism pathways displayed partition-specific distributions. Modules involved in melatonin synthesis, ClpB production, S-adenosylmethionine metabolism (SAM), propionate synthesis, and glutamate degradation were reduced in the *Bact-Firm* and *Prev* partitions. Conversely, serotonin metabolism, cortisol degradation, p-cresol production, and GABA metabolism modules were enriched in *Bact-Firm*. The *Prev* partition additionally showed increased quinolinic acid production and acetate degradation. Kynurenine pathway modules were also differentially represented (Supp. Fig. S16). Together, these findings show that the distinct mental health profiles across partitions are accompanied by differences in microbial neuroactive potential.

## 3 Discussion

Despite extensive efforts to characterize the determinants of human gut microbiome variation [9, 22–24], the ecological processes shaping microbiome configurations and their relationships with host health remain poorly understood. To address this gap, we developed a rule-based partitioning framework to stratify the *Le French Gut* population according to individual enterosignature profiles and explore associated functional and health-associated patterns. This stratification approach, designed to be applicable across cohorts, identified seven main partitions characterized by dominance or co-dominance of the ES-*Prevotella*, ES-*Firmicutes*, and ES-*Bacteroides* signatures and revealed functional gradients across the gut microbiome continuum. Patterns identified in the *Le French Gut* cohort were reproduced in an independent meta-cohort using the same partitioning approach. Integrating dietary, health, and fecal glycomic data further allowed us to explore the ecological context of these partitions and their potential relationships with host health.

A striking observation was the contrasting distribution of microbial functional potential related the three major hydrogen disposal pathways across partitions, reproduced across both cohorts. H_2_ is generated during carbohydrate fermentation, reaching up to *∼* 13 L per day in the human gut [25]. Its accumulation increases H_2_ partial pressure and inhibits fermentation unless it is removed by H_2_-consuming microorganisms including sulfate-reducing bacteria, methanogens and, to a lesser extent, acetogens [26]. In our dataset, these three pathways were distributed along the continuum of partitions. We therefore investigated the factors that might explain this variation and how they relate to host health.

*Prevotella*-enriched partitions showed a higher prevalence of modules of the sulfate reduction pathway, which can be energetically favored over methanogenesis when sulfate is available [26, 27]. Dietary and exploratory glycomic analyses point to two potential sulfur sources favoring this metabolism. The first is wine, which contributed to dietary sulfite exposure. The second is the more highly sulfated mucins observed in these configurations, from which sulfate could be released by the sulfatase activity reported in *Prevotella* species [28, 29] and then made available to sulfate-reducing partners, a role previously demonstrated for *Bacteroides* species [30–32].

*Prevotella*-enriched configurations have traditionally been associated with fiber-rich diets, particularly in non-Western populations [33], but their ecological determinants remain incompletely understood in industrialized populations [34–36]. Our study highlights sulfur and hydrogen metabolism as additional dimensions of *Prevotella*-enriched configurations. However, the identification, in our study, of two distinct *Prevotella* partition subpopulations differing in microbial diversity, without clear differences in the environmental or dietary factors assessed, underscores the need to better investigate their ecological determinants.

Conversely, methanogenesis modules were enriched in *Firmicutes*-associated partitions, together with butyrate-production potential and alpha diversity, consistent with previous reports [37, 38] and a fiber-supported fermentative ecosystem in which H_2_ production may sustain methanogenic activity. Longer transit times in these partitions, as suggested by lower Bristol Stool Scale scores, could favor the persistence of slow-growing microorganisms and contribute to the coexistence of greater richness and methanogenesis potential [38, 39]. Causality may also run in the opposite direction, since methane has itself been reported to slow intestinal transit [40]. *Firm-Prev* and *Prev-Firm* partitions particularly exhibited healthier lifestyle and dietary habits and reported lower perceived stress, less frequent digestive symptoms and lower psychotropic drug prescription rates.

*Bacteroides*-dominant partitions showed greater acetogenesis potential, whereas methanogenesis and sulfate-reduction potential were found limited. One possible explanation involves oxidative stress, as *Bact-Firm* exhibited increased genetic potential for oxidative stress responses, including superoxide dismutase and catalase modules, alongside reduced abundance of sialylated core 3 mucin structures, a glycomic alteration also reported in patients with Crohn’s disease [18]. Together, these observations suggest a possible role for local redox conditions in shaping hydrogen disposal pathways, as some acetogens have been reported to exhibit greater tolerance to oxidative stress [41].

Lower dietary fiber intake and greater mucin-degradation signature in *Bact-Firm* suggest a potential shift from dietary fiber toward mucin glycans as microbial substrates, consistent with evidence that fiber deprivation promotes microbial mucin utilization [19]. Both substrates can fuel fermentation and H_2_ production, linking substrate use to hydrogen metabolism [25, 32]. Greater processed foods consumption in *Bact-Firm* may also imply higher exposure to food additives, some of which have been linked to mucus barrier alterations and inflammation, potentially favoring mucin degradation [42].

Our functional findings provide an ecological perspective on this phenotype and suggest a hypothetical model in which altered hydrogen handling contributes to its development. Hydrogenotrophic acetogenesis requires higher H_2_ partial pressures than methanogenesis or sulfate reduction due to its lower thermodynamic efficiency [43]. This can lead to H_2_ accumulation, which may inhibit fermentative metabolism and reduce butyrate production [27, 44]. Together with mucin degradation, reduced butyrate availability could compromise epithelial barrier function [45], a feature implicated in systemic diseases [46]. The lower butyrate-production potential and increased oxidative stress signatures in *Bact-Firm* are compatible with this model. Although speculative, our findings position hydrogen metabolism as a potentially central, yet overlooked, link between gut microbial ecology and host health.

The poorer health profile of *Bact-Firm* partition also extended to psychological well-being, as indicated by higher perceived stress and more frequent psychotropic prescriptions, although prescription records do not confirm that the medications were taken and may relate to other indications. Enrichment of GABA and serotonin biosynthesis modules in *Bact-Firm* suggests that greater microbial genetic potential for producing these compounds does not necessarily correspond to better mental health [47, 48]. Conversely, reduced SAM biosynthesis potential offers another hypothesis to explore, given reported improvements in depressive symptoms following supplementation [49, 50]. The accompanying oxidative stress signatures raise the possibility that host low-grade inflammation and activation of the tryptophan-kynurenine pathway may contribute to the poorer mental health indicators associated with *Bact-Firm* [51].

Overall, the *Bact-Firm* partition was characterized by low microbial diversity and a less favorable health profile, including higher BMI, a greater burden of chronic disease, and more frequent digestive symptoms, resembling previous descriptions of the *Bacteroides* 2 enterotype [21, 52]. The stratification procedure also identified a *Bact* partition comprising 73 individuals, which was merged with *Bact-Firm* for statistical analyses because of its small size. The higher proportion of *Bact* individuals in the international meta-cohort suggests that this partition may represent a distinct subpopulation whose ecological and health characteristics warrant further investigation.

Beyond their ecological interpretation, our results illustrate the value of combining metagenomics with fecal mucin O-glycosylation profiling in a general population cohort, extending its application beyond inflammatory bowel disease [53]. This complementary approach provides a non-invasive biochemical perspective on mucus–microbiota interactions, with potential relevance to intestinal barrier function.

Several limitations should be acknowledged. First, the cross-sectional design precludes causal inference, and the proposed model should be viewed as a hypothesis-generating framework. Second, hydrogen metabolism, oxidative stress, and butyrate production were inferred from gene content. Genome-scale metabolic modeling could help refine the resulting hypotheses, while metatranscriptomic analyses could determine whether the inferred metabolic pathways are actively expressed [54]. Third, dietary data were self-reported and subject to recall bias. Fourth, residual confounding by medication, age, sex, BMI and overall diet cannot be excluded. Finally, our cohorts are drawn from specific populations and generalization to other settings remains to be tested.

In conclusion, we developed an enterosignature-based partitioning framework that adds an ecological perspective to gut microbiome investigation. By combining the quantitative resolution of continuous ES profiles with discrete, interpretable groups, this framework may facilitate inter-study comparisons and provide a practical basis for future clinical stratification. It could also support the design of stratified microbiome-directed interventions. One potential application is precision nutrition, where partition membership could help identify and prioritize small, manageable dietary or lifestyle adjustments aimed at modulating relevant microbial functions, particularly in the *Bact-Firm* partition. Such targeted strategies may be more acceptable and easier to sustain than broad behavioral changes [55]. Prospective intervention studies are needed to establish the clinical utility of these partitions and determine whether modifying the identified microbial functions improves host health.

## 4 Material & Methods

### 4.1 The *Le French Gut* project design

*Le French Gut – Le Microbiote Français* is an ongoing cross-sectional study initiated in September 2022 in metropolitan France. Participants were recruited from all metropolitan French regions on the LFG platform (https://le-french-gut.skezia.io) hosted by Skezia, where all data collection took place. The platform complies with French Health Data Hosting (HDS) regulations and the EU General Data Protection Regulation (GDPR). The primary objective of the study is to investigate associations between diet, health, lifestyle, and gut microbiota composition in adults (*≥* 18 years old). The study design and protocol have been previously published [13]. Eligible participants were adults who gave informed e-consent and had no history of colectomy, digestive stoma, recent colonoscopy (*<* 3 months), or recent antibiotic use (*<* 3 months). The study was approved by the “Comité de Protection des Personnes” (CPP Sud-Est IV, reference 21.00225.000006), authorized by the CNIL (*Commission Nationale de l’Informatique et des Libertés*) (ref: DR-2022-141), and registered under ID 2021-A01439-32.

### 4.2 Participants included in this study

This study focuses on the first 5,169 participants of the LFG project. To take part, volunteers had to answer a first-floor online questionnaire about their health, lifestyle, and dietary habits, and collect a compliant stool sample. Optional surveys were available, including the optional MS-Nutrition Food Frequency Questionnaire [56], which was completed by 1,696 participants.

### 4.3 Metagenomic datasets

#### 4.3.1 Le French Gut

Stool samples were received by the L’Assistance Publique - Ĥopitaux de Paris (APHP) laboratory in Bobigny. They were then collected and processed as previously described by Connan *et al.* [13]. DNA was purified from sample aliquots using the publicly available Epsilon pipeline [57].

Sequencing was performed using DNBSEQ technology (MGI Tech). Prior to sequencing, DNA degradation was checked using the DNA genomic 50kb kit on a Fragment Analyzer 5200 (Agilent) [58]. After fragmentation of 100 to 500 ng of DNA, libraries were constructed according to the instructions in the MGIEasy Universal DNA Library Prep Set, FS or Fast FS Library Prep Set (MGI), including end repair, A-tailing, adapter ligation, and selection of constructs by amplification. Barcoded libraries were circularized using the MGIEASY Circularization Kit. Quality controls were performed using Small Fragment kits on a Fragment Analyzer 5200 (Agilent Technologies), Quant-iT dsDNA and Qubit ssDNA Assay Kits (Thermo Fisher Scientific). Circularized library pools are constructed to optimize the yield of 20M paired-end reads (2 x 150) on a DNBseq G400 sequencer (MGI). Most steps are automated using Beckman Coulter and MGI pipetting handlers.

Remaining DNA and sample aliquots were stored at -70 °C in the MetaGenoPolis-SAMBO BRC (Biological Resources Center), part of INRAE (French National Research Institute for Agriculture, Food, and the Environment).

#### 4.3.2 International meta-cohort

We prioritized interpretability and biological relevance by using *Le French Gut* as the primary discovery cohort and the meta-cohort as an independent replication set. The meta-cohort includes samples from 8 public studies [20, 59–65] which were selected to mirror the LFG cohort in terms of metadata distribution [66] (Supp. Fig. S9). Minimal metadata for cohort inclusion included: age, sex, Body Mass Index (BMI) and health status. Their distribution was used for cohort selection in order to match the LFG distribution.

For each of the 8 studies, whole metagenome sequencing data was downloaded from the European Nucleotide Archive (ENA). Raw metadata were collected from ENA, NCBI, and supplementary materials of the original publications. Variables were standardized across studies, and a common set of minimal metadata was retained (age, sex, BMI, health status, country, study name).

To obtain the final meta-cohort (5,107 samples), samples with a value of “yes” in the “flag” column of the metadata table available at https://doi.org/10.57745/UPITJ0 were removed. Samples were flagged following several criteria: samples originating from longitudinal studies (time points beyond baseline, *i.e.*, t *>* 1) were excluded as well as samples with missing health status and current antibiotics use. Moreover, samples from public studies were retained only if their sequencing depth was *≥* 10M reads (single-end or paired-end), depending on the cohort, to ensure comparability with the sequencing depth range of the LFG cohort.

### 4.4 Metagenomic sample processing

Both cohorts were processed with the same tools and parameters, using the same pipeline with Meteor2 [67], primarily optimized for bacterial detection. Except for enterosignature calculations, all analyses were performed on sequencing data downsized to 20M single reads for the meta-cohort and 20M paired reads for the LFG cohort.

More precisely, for public cohorts originally sequenced in paired-end mode, only the forward (R1) reads were retained to ensure consistency with the single-end format of some cohorts. To reduce biases related to sequencing depth and ensure comparability with the LFG cohort, sequencing depths were constrained to a maximum twofold range. In the LFG cohort, sequencing depth ranged from 11M to 20M read pairs, with all samples downsized to a maximum of 20M read pairs. Accordingly, in the metacohort, sequencing depth ranged from 10M to 20M single-end reads after processing.

#### 4.4.1 Quality control

All DNA sequencing reads were quality trimmed and filtered from sequencing adapters using fastp [68] (v0.23.2). Remaining contamination by the host genome was removed by aligning the reads against the human reference genome T2T-CHM13v2.0 with Bowtie2 [69] (v2.5.1) and using at least 90 % nucleotide identity threshold for filtering, employing samtools [70] (v1.9).

Cross-sample contamination was checked with CroCoDeEL (v1.0.8) [71]. Samples were excluded if species added by contamination exceeded 12 % or if the contamination rate was *>* 1 % with additional species exceeding 10 %.

#### 4.4.2 Taxonomic profiling

For all samples, genes and microbial species were identified and quantified with Meteor2 (v2.0.18) [67] using human gut microbial gene catalog (IGC2, comprising 10.4 million genes), with GTDB R226 release for taxonomic annotation.

The human gut catalog was organized into 1990 Metagenomic Species Pangenomes (MSPs) [72] thanks to MSPminer [73]. An MSP is a cluster of co-abundant genes corresponding to a microbial species. MSP abundance was calculated as the mean abundance of its 100 marker genes (*i.e.*, species-specific core genes showing the strongest co-abundance). MSPs with fewer than 10 % of marker genes detected in a sample were assigned a null abundance.

MSP abundance tables for all samples were generated by Meteor2 with the command meteor profile -n coverage and read downsizing (rarefaction) to 20M paired reads per sample. This downsizing step was performed upstream before the mapping stage to equalize sequencing depth across samples before quantification, thereby controlling for unequal sequencing effort so that species-detection sensitivity and abundance estimates are comparable between samples. Profiling at the gene level involved a second normalization step relating to coverage. Each gene’s read count was converted into an estimate of coverage over the gene, correcting for gene length and for edge effects caused by read trimming during mapping. Overall, for each MSP, Meteor2 considers its core genes and checks their fraction having a non-zero (normalized) count in the sample. Ultimately, an MSP abundance equals to the mean of the (already-normalized) gene-level values across those detected core genes. The values in the MSP abundance table are coverage-normalized signature-gene abundances, *i.e.*, depth-of-coverage-like quantitative estimates, not proportions that sum to 1 across MSPs within a sample.

#### 4.4.3 Functional profiling

Samples were profiled at species, functional and strain level with Meteor2 (v2.0.18, normalization = coverage, references = human gut). Briefly, Meteor2 maps metagenomic reads on gene catalogues (clustered into Metagenomic Species (MSP) and functionally annotated) and further infers (i) species (MSP) abundance table, (ii) inter-sample genetic distance for each MSP (strain level) (iii) functional modules (KEGG, Gut Metabolic modules (GMM) and Gut-brain Modules (GBM)) abundance table. MSPs were taxonomically annotated using GTDB release r226.

A functional module is considered to be present in an MSP if at least 90 % of its components are found among the MSP genes. Module abundance was quantified by summing the abundances of all MSPs carrying the module in each sample.

### 4.5 Enterosignature calculation & Partitioning

Enterosignature (ES) composition was calculated for both the LFG cohort and the international meta-cohort by applying the five-ES model described in [10], using cvanmf v0.3.1 [74]. Briefly, after total-sum scaling of the genus-level abundance profiles, non-negative sample weights were estimated against the five fixed enterosignatures by multiplicative-update non-negative matrix factorization using the generalized Kullback–Leibler divergence and no regularization. The weights were then normalized to represent the relative contributions of the five enterosignatures. The cosine similarity score obtained using the five-ES model was 0.80, consistent with the scores reported in [10] and supporting the reusability of the model. Each sample was therefore described by proportions, summing to one, of five ES characterized by *Bacteroides*, *Firmicutes*, *Prevotella*, *Bifidobacterium*, and *Escherichia* [10].

Following [10], we define the *primary ES* as the most abundant ES in a sample. We also calculated ES diversity as the minimum number of ES required to account for at least 90 % of the cumulative relative contribution in a sample, after ordering the ES by decreasing relative abundance.

Using the normalized ES contributions, we then applied a rule-based partitioning procedure to stratify samples according to the relative dominance of their primary ES (parameter *α*) and the combined contribution of their primary and secondary ES (parameter *θ*). Each sample was assigned to one of three configurations as described below.

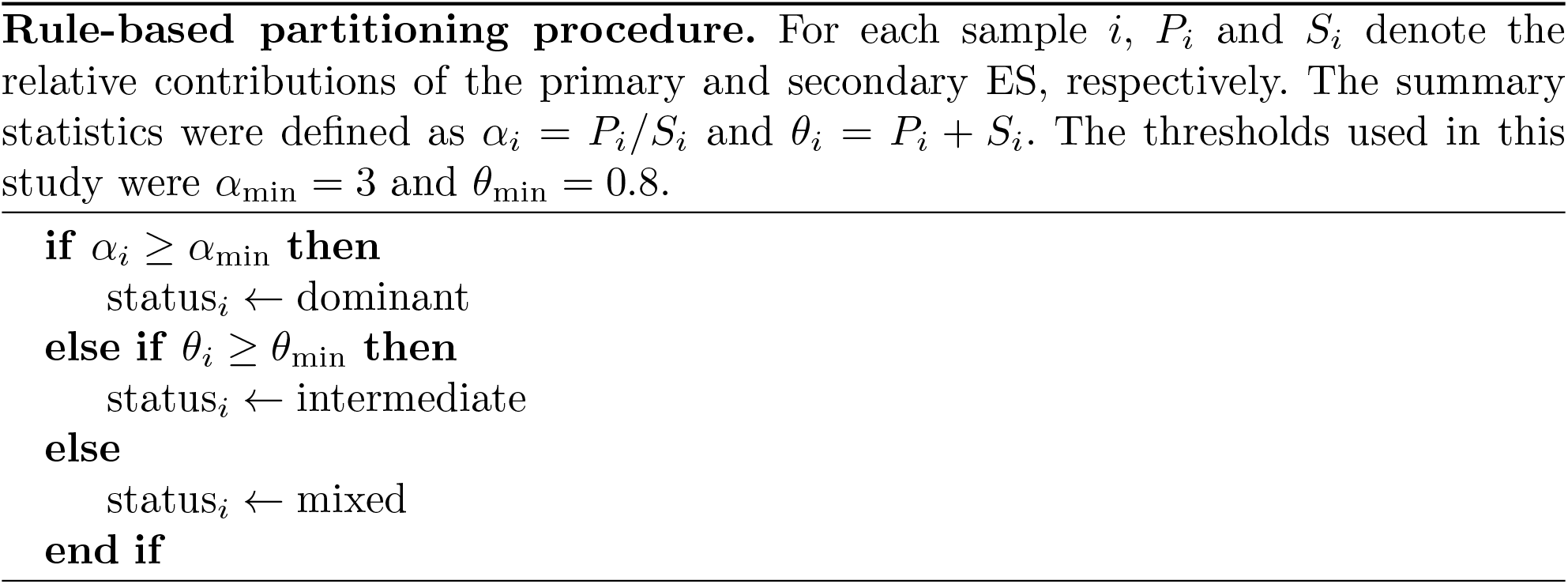

The threshold derivation and full partitioning procedure are detailed in the Supplementary Methods.

The final partition label was derived from the identities of the primary and secondary ES and the status assigned by Algorithm 1. A sample was assigned to a dominant partition when the ratio between its primary and secondary ES contributions was at least 3. Among the remaining samples, those for which the primary and secondary ES together accounted for at least 80 % of the ES composition were assigned to an ordered primary-secondary partition. All other samples were assigned to the *Mixed* partition.

Partitions containing fewer than 2 % of the samples were subsequently consolidated into a *Rare* category for population-level analyses, except for the *Bact* (n = 73) partition which was merged with *Bact-Firm*. All other rare partitions included less than 16 individuals. This procedure resulted in eight partitions in the LFG cohort. The same partitioning procedure generated nine partitions in the meta-cohort, with the *Bact* group here exceeding the 2 % threshold and the other partitions being identical to those from the LFG cohort. For consistency however, the *Bact* samples were again combined with those from *Bact-Firm* to ensure comparability with the LFG cohort.

For validation purposes, using the ES outputs of the LFG cohort, we also stratified the population into seven partitions with *k*-means and Partitioning Around Medoids (PAM) clustering algorithms (*k* = 7) and compared the cluster assignments to the partitions described above (Supp. Fig. S2c). The *k*-means clustering was applied on the ES proportions (normalized to 1 for each sample) and the PAM clustering was performed on the Bray-Curtis distances calculated from the previously mentioned table using the cluster R package (v2.1.8.2) [75].

To test the generalizability of our partitioning procedure, we applied it to the dataset from Frioux *et al.* [10], which contains individuals from 13 countries and of a wide age range including infants, leading to more diverse ES profiles than those obtained in the LFG cohort (Supp. Fig. S2d).

### 4.6 Statistical analyses on taxonomic profiles

Differential analysis on the abundance of metagenomic species across all partitions was done with the Kruskal-Wallis test. The tests were performed on the abundance data normalized with a clr transformation. The top 3 drivers per partition were identified after focusing on the three archetypal partitions *Bact-Firm*, *Firm* and *Prev*, and filtering for prevalent species (*i.e.* prevalence *≥* 30 %). MSP drivers were identified as species dominant in one partition compared to the two other ones. The ranking was obtained by ordering their mean Cliff’s Delta in absolute value. The detailed list is in the Supp. Table 1.

### 4.7 Statistical analyses on functional profiles

#### 4.7.1 Module prevalence

For prevalence analysis of functional modules, we focused on the three archetypal partitions: *Bact-Firm*, *Firm* and *Prev*. We calculated module prevalence in each partition and defined Δ as the difference in prevalence between one partition and the two other ones. Modules with Δ *≥* 10 % were retained for further analysis and referred as discriminant. All modules linked to H_2_ recycling (Fig. 2a) were found in this filtered list. The complete list is in Supp. Table 2.

#### 4.7.2 Module abundance

Differential analysis of the abundance of functional modules across all partitions was done with the Kruskal-Wallis test. A post-hoc Cliff’s delta test was then performed with the R package effsize (v0.8.1) to assess effect size. Modules presented in Fig. 2b were significant according to the corrected p-value (q-value *<* 0.05, see section 4.12) with at least one comparison showing an absolute value of Cliff’s Delta *>* 0.6. A selection of these is shown in Fig. 2b, and a list of all those that were significant can be found in the Supp. Table 3.

### 4.8 Mucin glycosylation analyses

We selected samples for experimental mucin analysis from the partitions *Bact-Firm* (n = 7), *Firm* (n = 7), and *Prev* (n = 12). Selected *Prev* samples included both high (n=6) and low (n=6) diversity samples following the observation of a large distribution range of Shannon diversity indices. The *Bact-Firm* group included 3 *Bact* samples.

Sample selection excluded individuals with chronic disease, and those taking dietary supplement consumption and medications other than painkillers or oral contraceptives. Participants with recurrent infections in the past year, or with antibiotic use or COVID-19 infection within the past 3 months were also excluded.

Samples were selected within each subgroup to ensure equal representation of females and males, and to achieve age balance, with a mean age of approximately 35-40 years. We also considered the consumption of foods associated with sulfite intake (*e.g.*, alcohol and nuts), preferentially selecting individuals with lower reported consumption to minimize potential dietary bias.

#### 4.8.1 Mucin purification

Stool samples were suspended in sodium chloride solution (0.2 mol/L) containing 0.02 mol/L sodium azide at 4°C. After homogenization, the samples were immediately centrifuged at 10 000 g for 30 min. The supernatant was dialyzed, freeze dried and mucins were solubilized in extraction buffer (containing 4 M guanidine chloride, 5 mM ethylenediaminetetraacetic acid (EDTA), 10 mM benzamidine, 5 mM N-ethylmaleimide, 0.1 mg/mL trypsin inhibitor, and 1 mM phenylmethane-sulfonyl fluoride), and then purified by isopycnic density-gradient centrifugation (Beckman Coulter LE80K ultracentrifuge; 70.1 Ti rotor, 417 600 g at 15°C for 72 h). Mucin-containing fractions were pooled, dialyzed, and lyophilized before use.

#### 4.8.2 Glycan release and permethylation

Mucins were then submitted to *β*-elimination under reductive conditions (0.1 M KOH, 1 M KBH4 for 24 h at 45°C). After co-evaporations with methanol, oligosaccharides were purified on a cation exchange resin column (Dowex 50 x 2, 200-400 mesh, H+ form). The oligosaccharide-alditol fractions were then permethylated, in their anhydrous form, in a solution containing 200 *µ*L dimethylsulfoxide, 300 *µ*L iodomethane, and 1 g NaOH, for 2 h, before adding 1 mL acetic acid (5 % (v/v)) to stop the reaction. After derivatization, the reaction products were dissolved in 200 *µ*L methanol and further purified on a C18 Sep-Pak column (Oasis HLB, Waters, Milford, MA, USA).

#### 4.8.3 MALDI-TOF mass spectrometry

Permethylated oligosaccharides were analyzed by matrix-assisted laser desorption ionization-time of flight (MALDI-TOF) mass spectrometry (MS) in positive ion reflective mode as [M+Na]+. Samples were dissolved in a methanol/water solvent (50:50) and coated on a MALDI target with a 2,5-dihydroxybenzoic acid matrix at a volume/volume dilution. The relative percent of each oligosaccharide was calculated, based on the integration of peaks on MS spectra.

### 4.9 Dietary assessment

#### 4.9.1 Dietary intake diversity

The Fruit and Vegetables Diversity index was estimated using self-reported habitual consumption from the year preceding participation. In the first-floor questionnaire, participants were asked which fruits and vegetables they consumed in the previous year. Each fruit or vegetables item was classified as 1 if consumed, and 0 otherwise. Each participant’s yearly Fruit and Vegetables Diversity index was determined by adding all fruits and vegetables reported as consumed, with higher scores indicating greater diversity of fruit and vegetables consumption.

#### 4.9.2 Food Frequency Questionnaire

Dietary intake was assessed using the optional MS-Nutrition Food Frequency Questionnaire (FFQ), a semi-quantitative, online 94-item instrument designed to estimate usual dietary intake over the previous month in French adults. The FFQ includes food items and portion sizes representative of foods commonly consumed in France. Each item corresponds to a group of related foods (*e.g.* raw vegetables, whole-grain starches), with portion size information collected for 50 items using standard units, household measures, or manufacturer-defined containers. Nutrient intakes were estimated using the French CIQUAL 2013 [76] food composition database, with item composition weighted according to consumption data from the INCA2 [77, 78] national dietary survey. Dietary sulfite intake from food and drink additives was estimated by combining published average sulfite concentrations for sulfite-containing foods to the individual daily consumption (g/day) of these food items [79, 80]. The MS-Nutrition FFQ has demonstrated acceptable validity and reproducibility for estimating daily energy intake (kcal/day), macronutrient intake, food group consumption, and dietary quality in the French adult population [56].

#### 4.9.3 aHEI-2010 index

Diet quality was assessed using the alternative Healthy Eating Index 2010 (aHEI-2010) [81, 82], which is based on 11 food and nutrient components with well-established associations with chronic disease and mortality. Each component was scored from 0 to 10, with higher scores reflecting higher intakes of vegetables, fruit, whole grains, nuts and legumes, long-chain omega-3 fatty acids, and polyunsaturated fatty acids, as well as lower intakes of sugar-sweetened beverages, red and processed meats, trans fatty acids, and sodium. For alcohol, moderate consumption received the highest score, whereas heavy consumption received the lowest score. Intermediate intakes were scored proportionally based on predefined cutoffs. Total aHEI-2010 scores ranged from 0 to 110, with higher scores indicating better overall dietary quality.

#### 4.9.4 Additional analyses

Spearman correlations were performed between MSP drivers’ abundance and dietary items’ consumption. P-values were adjusted for multiple testing (see section 4.12, Supp. Table 8).

#### 4.9.5 Water sanitary and quality checking

We obtained the sulfate concentration (mg/L) for each French commune (civil township) from the results of the governmental sanitary tap water control study in 2025 (https://www.data.gouv.fr/datasets/resultats-du-controle-sanitaire-de-leau-distribuee-commune-par-commune). We then averaged the measurements of each commune throughout the year 2025.

To proxy participants’ residential addresses, we used the IRIS code, which corresponds to a geographical area covering approximately 2,000 people (https://www.insee.fr/fr/information/7708995). As governmental water sanitary and quality checks data were accessible for each commune, the IRIS codes were matched, where possible, with commune codes.

### 4.10 Health & Lifestyle analysis

BMI categories presented in Fig. 6 follow the guidelines of the United States Centers for Disease Control and Prevention (CDC) [83]. In the data curation process, an unrealistic BMI value of 100 kg/m^2^ was considered as NA.

The DSFQ score is a composite score of digestive symptoms (abdominal pain, intestinal bloating, flatulence, and stomack gurgling) that was calculated as described by Azpiroz *et al.* [14]. When completing the questionnaire, individuals were asked to refer to the week preceding participation.

The PSS-4 score is a method to assess psychological stress according to Vallejo et al. [15]. It is calculated based on questions regarding self confidence, feeling good, feeling overwhelm, and feeling out of control. When completing the questionnaire, individuals were asked to refer to the month preceding participation.

Bristol Stool Scale scores were categorized as constipation (types 1–2), normal (types 3–5), and diarrhea (types 6–7), according to Blake *et al.* [84].

The Healthy Lifesyle Index (HLI) was constructed following the work of Sun *et al.* [85], and incorporates diet quality, physical activity, tobacco use, and alcohol consumption.

### 4.11 Drug prescription tracking

Data source and drug prescription tracking utilized the French National Heath Data System (*Système National des Données de Santé*, or SNDS) to access reimbursed health events for all participants in the LFG project. Each participant’s data were linked via their social security registration number (*numéro de carte vitale*, or NIR), enabling precise tracking of prescription records over time. The analysis was done on the 4,495 participants assigned to *Bact-Firm*, *Firm-Bact*, *Firm*, *Firm-Prev*, *Prev-Firm*, and *Prev* partitions that could be matched to the SNDS database with the NIR identifier. There were 136 unmatched participants who were uniformly distributed across the six partitions.

Prescription data for anxiolytic and antidepressant medications were surveyed for each participant, including the exact dates of reimbursement, and aggregated by partition. ATC classification codes were respectively N05B, and N06A [86]. We calculated the total number of prescriptions within three distinct time windows: two years, five years, and ten years prior to participation in the LFG project.

Prevalence rates (*i.e.*, prescriptions *>* 0) were determined by identifying the percentage of participants with at least one prescription for each medication class within the two-year, five-year and ten-year windows.

To assess potential differences in prescription frequency, participants with prescriptions were categorized as having a single prescription or more than one prescription, with the latter group further divided into tertiles based on prescription count (Supp. Figs. S13 and S15).

### 4.12 Other statistical analyses

Statistical and data analysis were performed in R v4.6.1 [87]. Graphics were generated with tidyverse (v2.0.0) [88], broom (v1.0.13) [89], ggpubr (v0.6.0) [90], patchwork (v1.2.0) [91], cowplot (v1.2.0) [92], ggtext (v0.1.2) [93], ggalluvial (v0.12.6) [94], ggpmisc (v1.0.0) [95], ggnewscale (v0.5.2) [96], and ggeffects (v2.3.2) [97].

All statistical tests were corrected using the Benjamini-Hochberg [98] procedure to control for multiple testing. The Bray-Curtis dissimilarities between samples were calculated for ES output and species abundances using vegan (v2.7-3) [99].

On the boxplot figures, results are shown using the compact letter display with multcompView (v0.1-11) [100] after Tukey’s HSD post-hoc test to assess pairwise significant differences. Two groups are significantly different if they do not share the same letter. In all boxplots, the box represents the interquartile (IQR) range and displays the median as a horizontal line, while the lower (respectively upper) whisker extends to the maximum (resp. minimum) between 1.5*∗*IQR and the smallest (resp. largest) value of the data.

Comparison analyses across partitions were performed using the Kruskal-Wallis and Chi2 tests for numerical and categorical variables, respectively. The variables shown in figures were all significant (q-value *<* 0.05). Extended lists of all significant tests can be found in the Supp. Tables.

Forest plots were based on multivariable regression models, using negative binomial regression and the continuous values of variables of interest (*e.g.* frequency of digestive symptoms, drug prescriptions rates). Age, sex, and BMI were selected as *a priori* covariates and the reference partition was set as *Firm*. Adjusted effect estimates were indicated on the figures with 95 % confidence intervals. The models were obtained using MASS (v7.3-66) [101].

Tweedie generalized linear model (GLM) regressions based on ES proportions were performed using statmod (v1.5.2) [102] to predict the functional modules’ abundances (Supp. Fig. S8). Tweedie GLM models were used with a log link function (power parameter *p* = 1.5) as it accommodates for the continuous, non-negative, and right-skewed distribution of the modules’ abundance data, including exact zeros. Each model included *ES-Bact*, *ES-Firm* and *ES-Prev* as fixed effects. To visualize the predicted effect of each predictor, we generated response curves by varying the focal predictor (one ES) across its observed range while holding the other two predictors constant at their sample means.

For strain analyses of the *Prev* partition subpopulations (Supp. Fig. S4), hierarchical clustering was done on the mutation rate and the hierarchical tree was cut at 0.015 genetic distance, approximately 97 % average nucleotide identity, which corresponds to subspecies threshold [103]. The heatmap was visualized using ComplexHeatmap (v2.20.0) [104], gridExtra (v2.3) [105], ggthemes (v5.2.0) [106], RColorBrewer (v1.1-3) [107].

## 5 Data availability

### 5.1 Le French Gut Cohort

To protect participant privacy and comply with the French Data protection authority (CNIL) authorization granted for the *Le French Gut* study, individual-level metagenomic data cannot be made openly available without access control. Metagenomic data have been deposited in the European Genome-phenome Archive (EGA) under accession number EGAXXXXX. Data can be made available upon request through the EGA data access mechanism. All requests will be reviewed by the Data Access Committee of the *Le French Gut* study. Proposals, researchers or institutions requesting data will be approved if they meet the standard criteria related to ethics, privacy and data protection regulations. If the collaboration is approved, a data access agreement will be required, and any necessary authorizations from the relevant administrative authorities may be needed. In compliance with existing regulations, no personally identifiable data, as well as SNDS data, will be accessible.

### 5.2 Meta-cohort

Processed metagenomic data, along with metadata information (sex, age, BMI, country and clinical status) and microbiome profiles for all participants of the Meta-cohort, are publicly available in an open access repository, at https://doi.org/10.57745/UPITJ0. Raw metagenomic samples are deposited in the European Nucleotide Archive (ENA) of the European Bioinformatics Institute (EBI) under accession numbers PRJEB39223, PRJEB37249, PRJEB11532, PRJNA834801, PRJDB4176, PRJNA319574, PRJNA530339, and PRJEB21528, and are publicly accessible.

## 6 Code availability

The data from the international meta-cohort is available at Entrepot Recherche Data Gouv https://doi.org/10.57745/UPITJ0. The R code developed for the analyses performed on both *Le French Gut* and the meta-cohort is available at https://forge.inrae.fr/french-gut/sola 2026 h2 metabo health.

## 7 Supplementary files

Supplementary methods and figures are available in supp material.pdf. Supplementary tables are available in supp tables.xlsx.

## 9 Author contributions

JDoŕe, DSE, RB, HB, MA, MB, CF, and PV conceived the project. CM, MG, BQ, ELC, FPO, FT, NP, CC, PW, PHO, GSK, CRM, CF, MB, and MS designed the methodology. SF and MS developed the software and GV, AH, SF, JD, and MS conducted the formal analyses. MB, CF, PV, CRB, JD, RL, and MS performed validation of the results. MB, CF, PV, and MS prepared the original draft of the manuscript. All authors contributed to the interpretation of the results, reviewed and edited the manuscript, and approved the final version.

## 11 Acknowledgments

We sincerely acknowledge Nessim Raouraoua for his help in developing bioinformatic tools facilitating analysis of mucin O-glycans.

The authors thank Mahendra Mariadassou, Simon Labarthe, Julien Tap, and Guilhem Sommeria-Klein for their contribution to scientific discussions and their guidance.

All authors extend their sincere thanks to the *Le French Gut* participants for their contribution to this citizen science initiative.

## 12 Consortium

Najate Achamrah^1,2,3^, Mathieu Almeida^4^, Anne-Sophie Alvarez^4^, Mourad Benallaoua^5^, Robert Benamouzig^5^, Magali Berland^4^, Oana Bernard^6^, Sylvie Binda^7^, Anne Blais^5^, Hervé Blottìere^4,8^, Elise Boreźee-Durant^4^, Ĺea Breton^9^, Alexandre Cavezza^4^, Benôıt Chassaing^10^, Lison Chevreau^4^, Möıse Cöeffier^1^, Chlóe Connan^4^, Anne-Marie Davila-Gay^11^, Lauren Demerville^12^, Śebastien Dime^4^, Kahina Djerrah^5^, Michel Dojat^13^, Jöel Doŕe^4,14^, Assia Dreux^15^, Inès Drouard^4^, Olivier Durlach^16^, Erik Eckhardt^17^, Alexandre Famechon^4^, Etienne Formstecher^18^, Cĺemence Frioux^4,19^, Śebastien Fromentin^4^, Nathalie Galleron^4^, Rozenn Gazan^20^, Amine Ghozlane^21^, Marine Gilles^4^, Oscar Gitton-Quent^4^, Lindsay Goulet^4^, Pedro H. Oliveira^22^, Anne Hiol^4^, Esra Ilhan^14^, Evelyne Jouvin-Marche^23^, Simon Labarthe^24^, Nicolas Lapaque^4,14^, Milan Lazarevic^12^, Emmanuelle Le Chatelier^4^, Marion Leclerc^4,25^, Julie L^e-Hoang^4^, Patricia Lepage^14^, Emmanuelle Maguin^14^, Matthieu Maillot^20^, Claudine Manach^26^, Emile Mardoc^4^, Mahendra Mariadassou^27^, Elliot Mathieu^4^, Nicolas Maziers^4,28^, Idir Mazouzi^5^, Juliette Meyer^21^, Béńedicte Monnerie^4^, Christian Morabito^4^, Christine Morand^26^, Julie-Anne Nazarre^29^, Sophie Nicklaus^30^, Anne-Sophie Nyob^31^, Florian Plaza Oñate^4^, Nicolas Pons^4^, Benôıt Quinquis^4^, Xavier Raffoux^4^, Etienne Ruppé^32,33^, Anne Rutigliano^31^, Jean-Marc Sabaté^5^, Jacques Sainte-Marie^34^, Mathilde Sola^4,19^, Guilhem Sommeria-Klein^24^, Manon Sudrie^4^, Julien Tap^14^, Florence Thirion^4^, Vincent Thomas^35^, Patrick Trieu-Cuot^36^, Karine Valeille^4^, Charline Vasseur^4^, Patrick Veiga^4,14^, Florent Vieux^20^, Giacomo Vitali^4^, Fabien Wuestenberghs^5^

^1^Univ. Rouen Normandie, Inserm, ADEN UMR1073, CHU Rouen, CIC-CRB 1404, Department of Nutrition, F-76000, Rouen, France

^2^Univ. Rouen Normandie, Institute for Research and Innovation in Biomedicine (IRIB), 76000, Rouen, France

^3^Department of Nutrition, CHU Rouen, 76000, Rouen, France

^4^Université Paris-Saclay, INRAE, MetaGenoPolis, 78350, Jouy-en-Josas, France

^5^Department of Gastroenterology, Avicenne Hospital, APHP, Université Paris Nord - La Sorbonne, Bobigny, France

^6^Biocodex, Gentilly, France

^7^Rosell Institute for Microbiome and Probiotics, Montreal, Canada

^8^Nantes Université, INRAE, UMR1280, PhAN, Nantes, France

^9^INSERM, Paris, France

^10^Microbiome-Host Interactions, INSERM U1306, CNRS UMR6047, Institut Pasteur, Université Paris Cité, Paris, France

^11^Université Paris-Saclay, AgroParisTech, INRAE, UMR PNCA, Palaiseau, France

^12^APHP, Paris, France

^13^Université Grenoble Alpes, Inserm U1216, Grenoble Institut Neurosciences, Inria, Grenoble, France

^14^Université Paris-Saclay, INRAE, MICALIS, 78350, Jouy-en-Josas, France

^15^Greentech, Saint-Beauzire, France

^16^Institut du Vieillissement, Hospices Civils de Lyon, Lyon, Auvergne-Rĥone-Alpes, France

^17^DSM-Firmenich, Houdan, France

^18^GMT Science, Paris, France

^19^Inria, University of Bordeaux, INRAE, 33400 Talence, France

^20^MS-Nutrition, Marseille, France

^21^Institut Pasteur, Université Paris Cité, Bioinformatics and Biostatistics Hub, F-75015, Paris, France

^22^Génomique Métabolique, Genoscope, Institut Fraņcois Jacob, CEA, CNRS, Université d’Évry, Université Paris-Saclay, Evry, France

^23^Institut Physiopathologie, Métabolisme, Nutrition (PMN) Inserm, Paris, France

^24^INRAE, BIOGECO, Univ. Bordeaux, Cestas, France

^25^Université Clermont Auvergne, INRAE, MEDIS, Clermont, France

^26^Université Clermont Auvergne, INRAE, UNH, F-63000 Clermont-Ferrand, France

^27^Université Paris-Saclay, INRAE, MaIAGE, 78350, Jouy-en-Josas, France

^28^Hospital Center Sud Francilien, Intensive Care Unit, Corbeil-Essonnes, France

^29^CarMeN Laboratory, Université Claude Bernard Lyon 1, INSERM, INRAE, Pierre-Bénite, France

^30^INRAE, Paris, France

^31^ANJAC, Paris, France

^32^Université Paris Cité and Université Sorbonne Paris Nord, Inserm, IAME, F-75018 Paris, France

^33^AP-HP, Ĥopital Bichat-Claude Bernard, Laboratoire de Bactériologie, F-75018 Paris, France

^34^Inria Paris, Team Ange, Paris, France

^35^Danone, Gif-sur-Yvette, France

^36^Institut Pasteur, Université Paris Cité, Unité de Biologie des Bactéries Pathogènes ‘a Gram-positif, Paris, France

