## Supplementary material for "Gut microbiome stratification reveals functional metabolic signatures linked to host health": Supp. Fig.

Mathilde Sola[1,2,*], Anne Hiol[1], Giacomo Vitali[1], Sébastien Fromentin[1], Marine Gilles[1], Emmanuelle Le Chatelier[1], Christian Morabito[1], Florian Plaza Oñate[1], Nicolas Pons[1], Benoît Quinquis[1], Florence Thirion[1], Jordan Denis[3], Renaud Léonard[3], Corinne Cruaud[4], Patrick Wincker[5], Pedro H. Oliveira[5], Guilhem Sommeria-Klein[2], Le French Gut Consortium, Catherine Robbe Masselot[3], Mathieu Almeida[1], Hervé Blottière[1,6], Joël Doré[1,7], Dusko S. Ehrlich[1], Robert Benamouzig[8,+], Clémence Frioux[2,1,+], Magali Berland[1,+], Patrick Veiga[1,7,+]

[1]Université Paris-Saclay, INRAE, MGP, 78350, Jouy-en-Josas, France

[2]Inria, Univ. Bordeaux, INRAE, 33400, Talence, France

[3]Université de Lille, CNRS, UMR 8576 - UGSF, Lille, France

[4]Genoscope, Institut François Jacob, CEA, Université Paris-Saclay, Evry, France

[5]Génomique Métabolique, Genoscope, Institut François Jacob, CEA, CNRS, Université Évry, Université Paris-Saclay, Évry, France

[6]Nantes Université, INRAE, UMR 1280, PhAN, Nantes, France

[7]Université Paris-Saclay, INRAE, MICALIS, 78350, Jouy-en-Josas, France

[8]Assistance Publique - Hôpitaux de Paris, Université Paris Nord - La Sorbonne, Bobigny, France

* Corresponding author, + co-senior last authors

### 1 Supplementary Methods

#### 1.1 Stratification of the LFG population

**Rationale** One objective of this study was to stratify the *Le French Gut* population into groups with more homogeneous enterosignature (ES) compositions. Given that variation in gut microbiome composition appears to follow a continuum rather than forming discrete clusters [3], we adopted a rule-based partitioning strategy that facilitates comparisons across the microbiome landscape without assuming the existence of naturally discrete groups.

Grouping individuals solely according to their primary ES can result in heterogeneous groups. For instance, the subpopulation with ES-*Firmicutes* as its primary ES includes both individuals in whom this ES is strongly dominant relative to the secondary ES and individuals exhibiting co-dominance with ES-*Bacteroides* or ES-*Prevotella*. Similarly, *Individual-1* (*ES-Prev* = 0.75, *ES-Firm* = 0.15, *ES-Bact* = 0.10) and *Individual-2* (*ES-Prev* = 0.45, *ES-Firm* = 0.43, *ES-Bact* = 0.12) share the same primary ES (*ES-Prev*) but exhibit markedly different ES compositions.

We therefore sought to stratify the population while distinguishing profiles characterized by the relative dominance of a single ES, the joint contribution of two predominant ES, or substantial contributions from more than two ES. In the LFG dataset, the primary ES was predominantly ES-*Firmicutes*, ES-*Bacteroides*, or ES-*Prevotella*. This approach distinguishes individuals with different ES compositions even when they share the same primary ES.

**Method** We developed a rule-based algorithm with a small number of parameters and constraints. The assignment rules were not calibrated to the distribution of individuals in the LFG dataset. The algorithm was designed to be transferable to other datasets analyzed using the same ES model, including populations whose ES profiles may differ from those commonly observed in large-scale studies of healthy Western adults (e.g., children, non-Western populations, or individuals with a specific disease).

Let $X \in \mathbb{R}_+^{G \times N}$ denote the matrix of genus-level relative abundances, with genera as rows and samples as columns, and let $W \in \mathbb{R}_+^{G \times K}$ denote the fixed matrix containing the $K$ enterosignatures. The matrix of non-negative sample weights, $H \in \mathbb{R}_+^{K \times N}$, was estimated by minimizing the generalized Kullback-Leibler (KL) divergence between the observed taxonomic profiles and their reconstruction with `cvanmf` v0.3.1 [4]:

$$\widehat{H} = \arg\min_{H \geq 0} D_{\mathrm{KL}}(X \parallel WH). \tag{1}$$

The optimization was performed using the multiplicative-update algorithm implemented in scikit-learn's `non_negative_factorization` function, with the ES matrix held fixed (`solver="mu"`, `update_H=False`, and `beta_loss="kullback-leibler"`). No regularization was applied. In the notation used here, $h_{ik}$ denotes the estimated weight of ES $k$ for sample $i$.

The relative contribution of ES $k$ to sample $i$ was calculated by normalizing the estimated weights:

$$p_{ik} = \frac{h_{ik}}{\sum_{\ell=1}^{K} h_{i\ell}}, \qquad \sum_{k=1}^{K} p_{ik} = 1. \tag{2}$$

For each sample, the normalized ES contributions were ranked in decreasing order,

$$p_{i(1)} \geq p_{i(2)} \geq \cdots \geq p_{i(K)}. \tag{3}$$

We defined $P_i$ and $S_i$ as the relative contributions of the primary and secondary ES, respectively, and $R_i$ as the summed contribution of the remaining $K - 2$ ES:

$$P_i = p_{i(1)}, \qquad S_i = p_{i(2)}, \qquad R_i = 1 - P_i - S_i = \sum_{j=3}^{K} p_{i(j)}. \tag{4}$$

A uniform ES composition was used as a mathematical reference profile:

$$\mathbf{p}_{\text{uniform}} = \left(\frac{1}{K}, \frac{1}{K}, \ldots, \frac{1}{K}\right). \tag{5}$$

For each sample, we calculated two summary statistics:

$$\alpha_i = \frac{P_i}{S_i}, \qquad \theta_i = P_i + S_i. \tag{6}$$

The ratio $\alpha_i$ quantifies the relative dominance of the primary ES over the secondary ES. A larger value indicates stronger relative dominance by a single ES. Importantly, this criterion captures relative rather than absolute dominance and can therefore identify both samples with a large value of $P_i$ and samples with a moderate value of $P_i$ but a small value of $S_i$.

The sum $\theta_i$ represents the proportion of the ES composition accounted for by the two predominant ES. A large value indicates that the sample can be mainly described by its primary and secondary ES.

Each sample was assigned to one of three statuses using the algorithm below. For the ES model used in this study, the thresholds were set to $\alpha_{\min} = 3$ and $\theta_{\min} = 0.8$, as described in the following section.

**Algorithm 1 Partitioning algorithm based on ES proportions.** For each sample, $\alpha_i$ and $\theta_i$ are calculated according to Eq. 6. The thresholds used in this study were $\alpha_{\min} = 3$ and $\theta_{\min} = 0.8$.

**if** $\alpha_i \geq \alpha_{\min}$ **then**
  $\text{status}_i \leftarrow$ dominant
**else if** $\theta_i \geq \theta_{\min}$ **then**
  $\text{status}_i \leftarrow$ intermediate
**else**
  $\text{status}_i \leftarrow$ mixed
**end if**

The final partition label was derived from the identities of the primary and secondary ES and the status assigned by Algorithm 1. For samples with a dominant status, the partition was named after the primary ES. For samples with an intermediate status, the partition was named after the ordered pair formed by the primary and secondary ES. Samples with a mixed status were assigned to the common *Mixed* partition.

For example:

- For

$$\text{ID1} = (\textit{ES-Prev} = 0.65,\ \textit{ES-Firm} = 0.20,\ \textit{ES-Bact} = 0.05,\ \textit{ES-Bifi} = 0.05,\ \textit{ES-Esch} = 0.05),$$

  we obtain $\alpha_1 = 0.65/0.20 = 3.25$. Because $\alpha_1 \geq 3$, ID1 is assigned a dominant status. As its primary ES is *ES-Prev*, its final partition is *Prev*.

- For

$$\text{ID2} = (\textit{ES-Prev} = 0.05,\ \textit{ES-Firm} = 0.55,\ \textit{ES-Bact} = 0.30,\ \textit{ES-Bifi} = 0.05,\ \textit{ES-Esch} = 0.05),$$

  we obtain $\alpha_2 = 0.55/0.30 = 1.83$ and $\theta_2 = 0.55 + 0.30 = 0.85$. Because $\alpha_2 < 3$ and $\theta_2 \geq 0.8$, ID2 is assigned an intermediate status. As its primary and secondary ES are *ES-Firm* and *ES-Bact*, respectively, its final partition is *Firm-Bact*.

- For

$$\text{ID3} = (\textit{ES-Prev} = 0.30,\ \textit{ES-Firm} = 0.25,\ \textit{ES-Bact} = 0.22,\ \textit{ES-Bifi} = 0.15,\ \textit{ES-Esch} = 0.08),$$

  we obtain $\alpha_3 = 0.30/0.25 = 1.20$ and $\theta_3 = 0.30+0.25 = 0.55$. Because neither condition is satisfied, ID3 is assigned to the *Mixed* partition. In this profile, the primary ES does not dominate strongly relative to the secondary ES, and the two predominant ES do not account for a sufficiently large proportion of the complete profile.

For a model containing $K$ ES, the algorithm can generate up to

$$K + K(K-1) + 1 = K^2 + 1 \tag{7}$$

partitions: $K$ dominant partitions, $K(K-1)$ ordered primary–secondary partitions, and one *Mixed* partition. With $K = 5$, the maximum number of partitions is therefore 26. This calculation treats reversed pairs, such as *Firm-Bact* and *Bact-Firm*, as distinct partitions because the first and second ES correspond to the primary and secondary ES, respectively.

For the population-level analyses performed in this study, we applied an additional dataset-specific consolidation rule: partitions containing fewer than 2 % of the samples were grouped into a *Rare* category. This rule was not part of the general assignment algorithm and may be adapted according to the sample size and analysis plan.

**Choice of parameter values** The thresholds $\alpha_{\min}$ and $\theta_{\min}$ were derived from the uniform reference composition defined in Eq. 5. For a profile containing $K$ ES, the contribution of each ES in this reference composition is $1/K$. We considered an ES to be enriched relative to this reference when its normalized contribution exceeded $1/K$. Here, "enriched" refers only to representation above the uniform reference value and does not imply statistical significance.

To define a profile characterized by two predominant ES, we imposed two criteria. First, we required the contribution of the secondary ES to reach or exceed the uniform reference value, *i.e.* being enriched:

$$S_i \geq \frac{1}{K}. \tag{8}$$

Second, we required the combined residual contribution $R_i$ of the remaining $K-2$ ES not to exceed the contribution of a single ES in the uniform reference composition:

$$R_i \leq \frac{1}{K}. \tag{9}$$

This criterion limits the total contribution not accounted for by the primary and secondary ES. It also implies that the tertiary ES does not exceed the uniform reference value, *i.e.* is not enriched, because $p_{i(3)} \leq R_i \leq 1/K$.

$$\theta_i \geq 1 - \frac{1}{K}. \tag{10}$$

We therefore defined the minimum threshold for the combined contribution of the primary and secondary ES as

$$\theta_{\min} = 1 - \frac{1}{K} = \frac{K-1}{K}. \tag{11}$$

Second, for a profile to be described by two predominant ES, we required the contribution of the secondary ES to reach the uniform reference value:

$$S_i > \frac{1}{K}. \tag{12}$$

From the definitions of $\alpha_i$ and $\theta_i$ in Eq. 6, we have

$$P_i = \alpha_i S_i \qquad \text{and} \qquad \theta_i = P_i + S_i = S_i(\alpha_i + 1). \tag{13}$$

Therefore,

$$S_i = \frac{\theta_i}{\alpha_i + 1}. \tag{14}$$

To define the boundary between relatively dominant and two-ES profiles, we set $S_i = 1/K$ when $\theta_i = \theta_{\min}$ and $\alpha_i = \alpha_{\min}$. Substituting these values into Eq. 14 gives

$$\frac{1}{K} = \frac{\theta_{\min}}{\alpha_{\min} + 1}, \tag{15}$$

and therefore

$$\alpha_{\min} = K\theta_{\min} - 1. \tag{16}$$

Using $\theta_{\min} = \frac{K-1}{K}$ from Eq. 11, we obtain

$$\alpha_{\min} = K\left(\frac{K-1}{K}\right) - 1 = K - 2. \tag{17}$$

In this study, we reapplied the five-ES model described by Frioux *et al.* [5], for which $K = 5$. The uniform reference composition was therefore

$$\mathbf{p}_{\text{uniform}} = \left(\frac{1}{5}, \frac{1}{5}, \frac{1}{5}, \frac{1}{5}, \frac{1}{5}\right), \tag{18}$$

and the resulting parameter values were

$$\theta_{\min} = \frac{K-1}{K} = \frac{4}{5} = 0.8, \qquad \alpha_{\min} = K - 2 = 3. \tag{19}$$

Because the algorithm first assigns samples satisfying $\alpha_i \geq \alpha_{\min}$ to a dominant partition, samples assigned an intermediate status satisfy

$$\alpha_i < 3 \qquad \text{and} \qquad \theta_i \geq 0.8. \tag{20}$$

These conditions ensure that the primary and secondary ES together account for at least 80% of the profile and that the contribution of the secondary ES is greater than 20%. At the boundary $\alpha_i = 3$ and $\theta_i = 0.8$, the secondary ES contribution is exactly 20%, but the sample is assigned to a dominant partition because the dominant condition is evaluated first.

Although the thresholds are expressed as functions of $K$, their suitability may depend on the number and composition of the ES included in the model. The thresholds derived here were applied to the five-ES model and may require validation before being used with other values of $K$.

**Application to the LFG cohort and reapplicability in other datasets** Applications of this stratification algorithm to the LFG data and the meta-cohort are described in the main results. Supp. Fig. S1C illustrates the increased homogeneity of individuals within partitions with respect to their grouping based on primary ES.

The algorithm can theoretically be applied to any dataset, when provided its ES output. The rules we have defined and the values of the two parameters $(\alpha, \theta)$ we have used are valid if the goal is to stratify a population according to its main possible configurations, focusing on samples that are strongly dominated by one ES or samples that are mainly described by two ES. If the goal is to further stratify the population by focusing on tertiary ES, additional hypotheses should be included and the algorithm should be adapted accordingly.

As a complementary example, we applied the partitioning strategy to the dataset associated to the 5-ES model from Frioux *et al.* [5]. The partitioning generated twelve partitions including some that were absent or considered in the *Rare* partition in the LFG and meta-cohort: *Bact-Bifi*, *Bifi*, and *Esch.* Results are illustrated in Supp. Fig. S2d.

### 2 Supplementary Figures

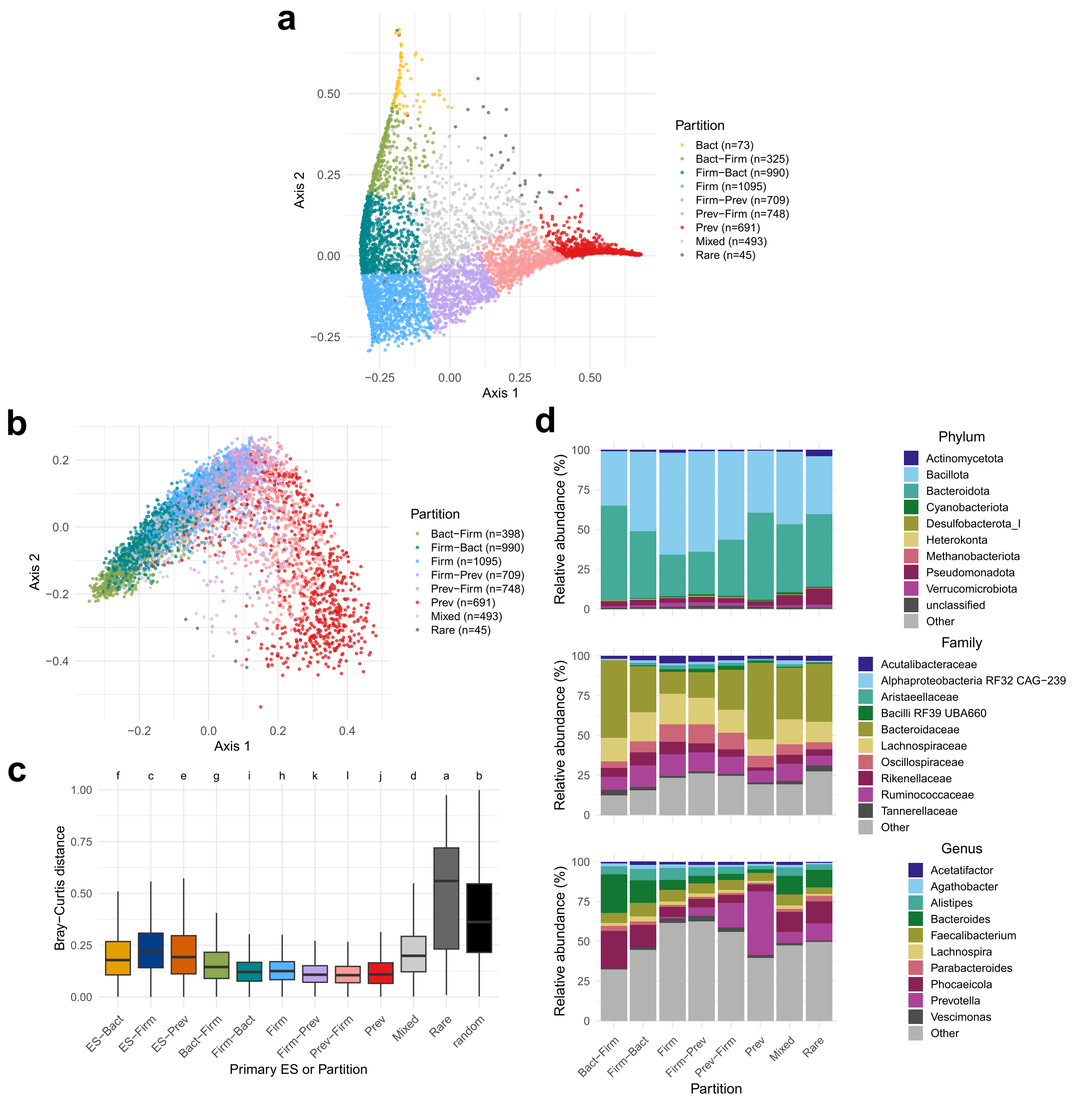


Figure S1: **Characteristics of the 7 partitions identified in the LFG cohort** a) PCoA of Bray-Curtis dissimilarities based on the ES composition of the LFG samples with the *Bact* partition distinguished from the Bact-Firm one. b) PCoA of Bray-Curtis dissimilarities based on the metagenomic species abundance data of the LFG samples. Each sample is colored according to its partition assignment. c) Intra-group Bray-Curtis dissimilarities in the LFG cohort. Boxplots indicate the range of distances calculated between pairs of samples from the same primary ES or partition. The random group corresponds to 5 random subsamplings of 500 individuals each, drawn from the entire LFG population. Letters correspond to the result of Tukey's HSD post-hoc test to assess pairwise significant differences. Different letters indicate that groups are significantly different. d) Relative abundances of the 10 most abundant phyla, families, and genera across partitions. The Other label includes all remaining taxa.

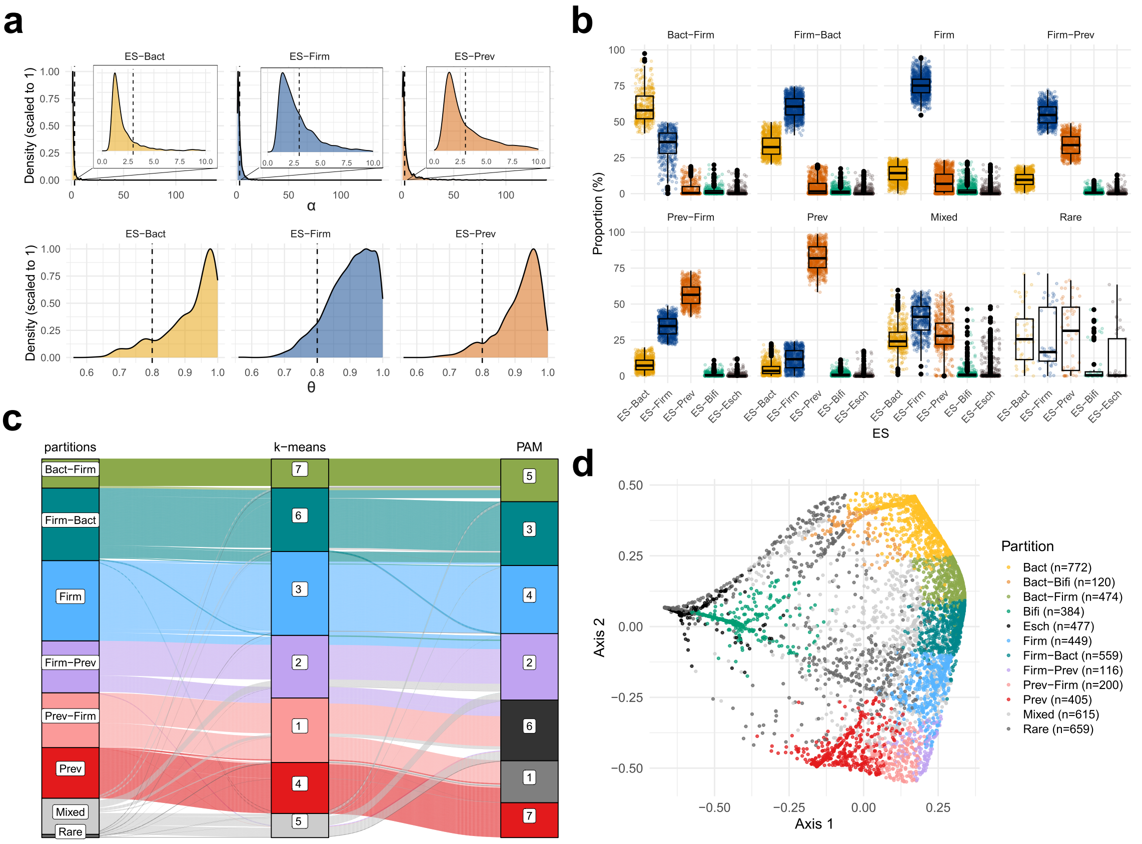


Figure S2: **Validation of the partitioning against clustering and application of the algorithm to the data of [5]** a) Distribution of the values of $\alpha$ and $\theta$ parameters involved in the partitioning algorithm across samples grouped by their primary ES. Values of $\alpha_{min}$ and $\theta_{min}$ selected in this study are illustrated with the vertical dashed lines. b) Distribution of enterosignature relative proportions across partitions in the LFG cohort. Boxplots indicate the range of ES proportions for each partition. c) Correspondence of samples between partitions and results from clustering methods (k-means and PAM) with $k = 7$. d) PCoA of Bray-Curtis dissimilarities based on the ES composition of the samples from Frioux et al. 2023 and colored by partition after applying the same algorithm as for the LFG cohort.

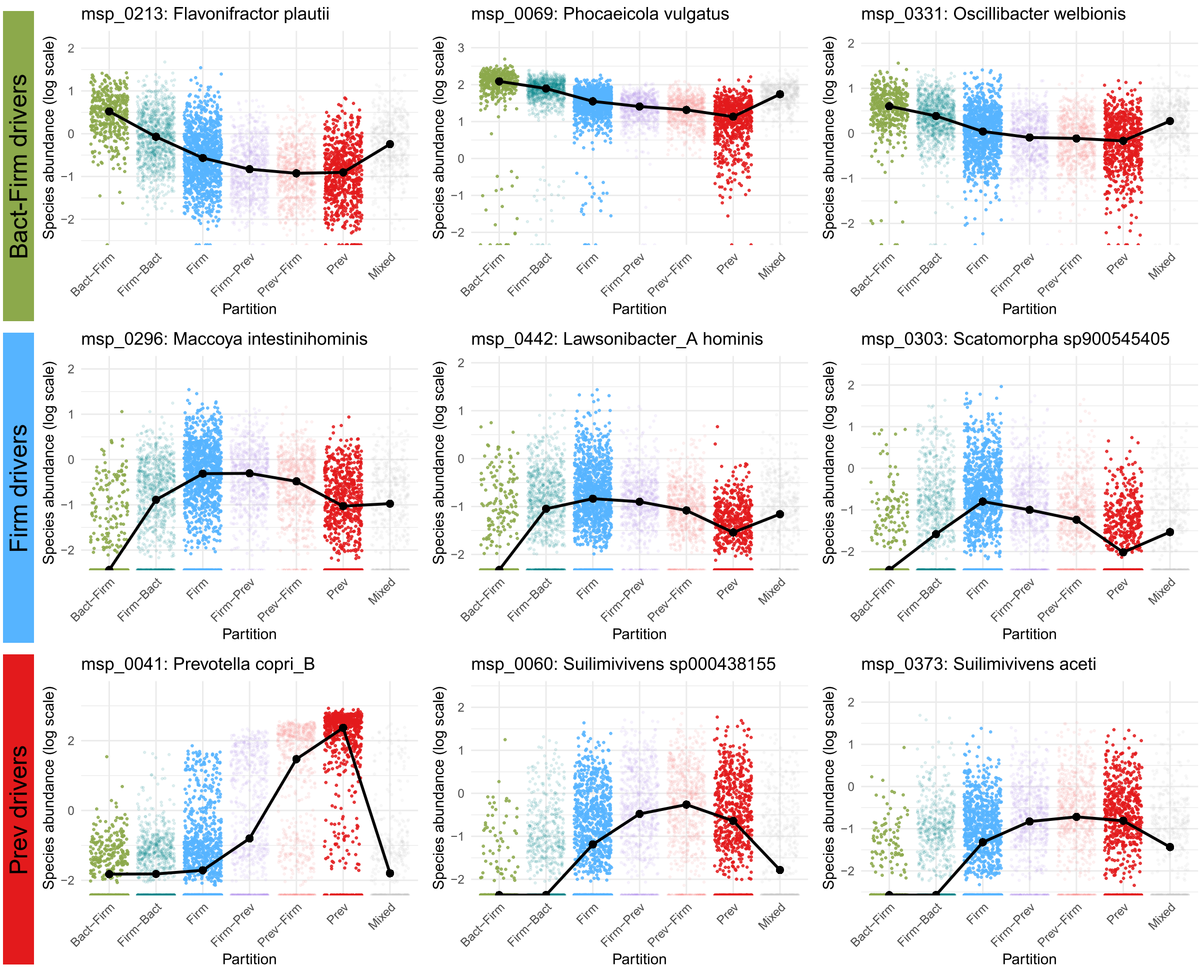


Figure S3: **Abundance of partitions species drivers across all individuals.** The top 3 drivers per partition were identified thanks to a differential abundance analysis on the metagenomic abundance data with Kruskal-Wallis tests across the three partitions *Bact-Firm* (top), *Firm*(middle) and *Prev* (bottom). Species drivers were identified as species dominant in one partition compared to the two other ones. The ranking was obtained by ordering their mean cliff's delta in absolute value. Absolute values of cliff's delta for each comparison is indicated on each plot. Identified drivers are: msp_0213: *Flavonifractor plautii*, msp_0069: *Phocaeicola vulgatus*, msp_0046: *Oscillobacter welbionis*, msp_0331: *Maccoya intestinihominis*, msp_0303: *Scatomorpha sp900545405*, msp_0442: *Lawsonibacter A hominis*, msp_0041: *Prevotella copri B*, msp_0373: *Suilimivivens aceti*, msp_0060: *Suilimivivens sp000438155*).

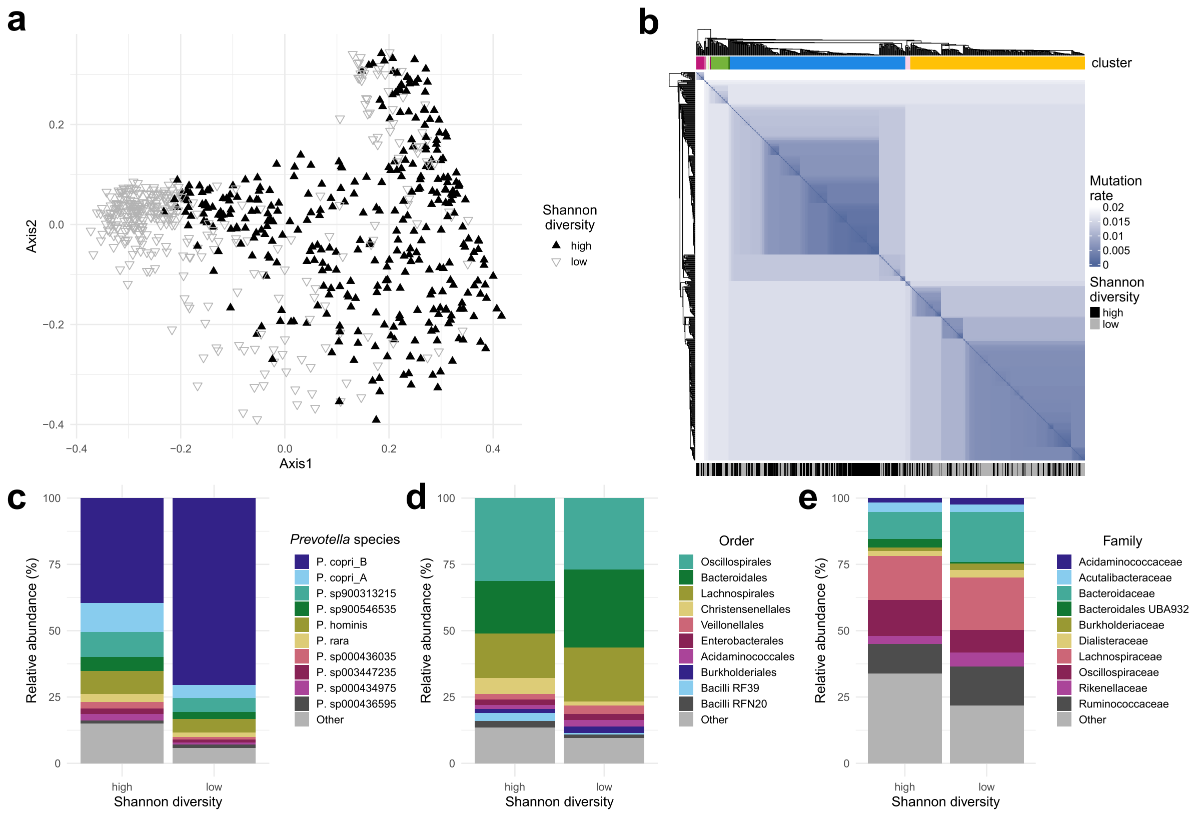


Figure S4: ***Prev* subpopulations defined by alpha-diversity and their characteritics** a) PCoA of Bray-Curtis dissimilarities based on the metagenomic species abundances of samples from the *Prev* partition. Points are distinguished according to whether the Shannon diversity value of the sample is above or below the median. b) Heatmap of pairwise genetic distances between *Prevotella copri B.* strains from different individuals. Clusters were defined by cutting the tree at a distance threshold of 0.015, corresponding to 97% average nucleotide identity (ANI). The bar on the top corresponds to this clustering. The bar on the bottom indicates whether the sample belongs to the high or low group according to Shannon diversity. c) Relative abundances of the 10 most abundant species among the *Prevotella* genus, for each subgroup established according to Shannon diversity. d) Relative abundances of the 10 most abundant orders, for each subgroup established according to Shannon diversity. e) Relative abundances of the 10 most abundant families, for each subgroup established according to Shannon diversity.

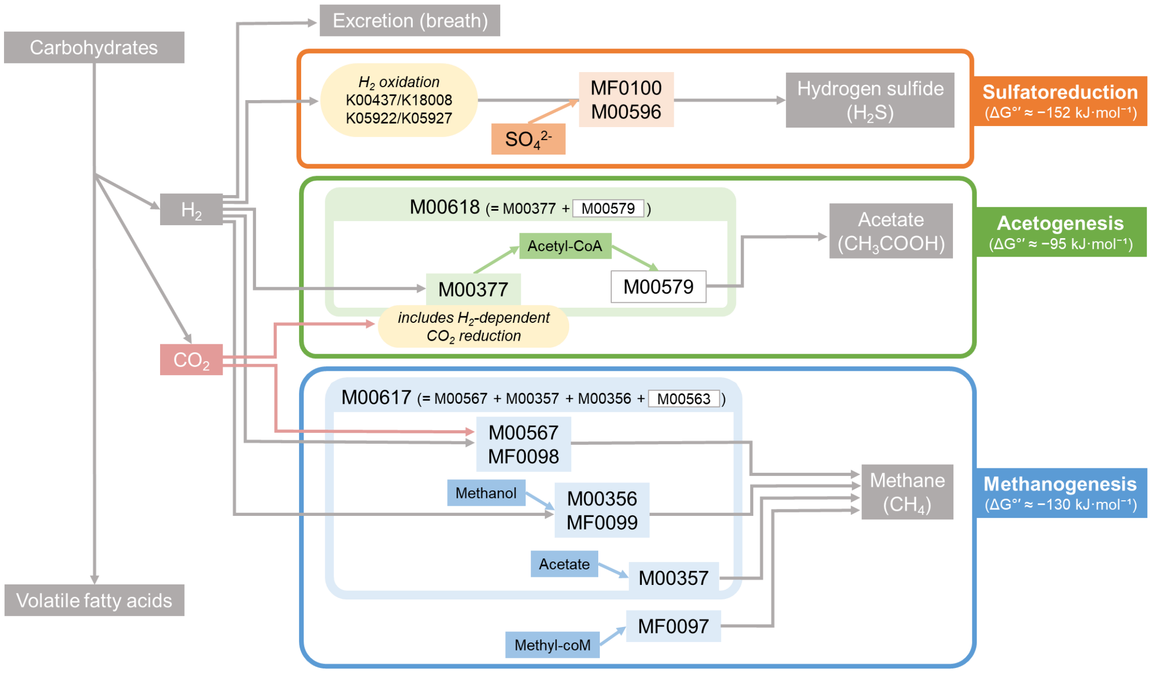


Figure S5: **Diagram of dihydrogen disposal in the human gut, adapted from Gibson *et al.* [6]** The 10 functional modules identified in Fig. 2a are flagged on the corresponding routes, the modules in white were not identified in the occurrence analysis shown in Fig. 2a. The standard Gibbs free energy ($\Delta G'$, *i.e.* the maximum amount of usable work a reaction can perform under biochemical standard conditions) is indicated for each route (Smith et al., 2019) [7]. A negative value of $\Delta G'$ means the reaction is spontaneous. Sulfatoreduction has the lowest value and therefore is thermodynamically the most favorable route among the three.

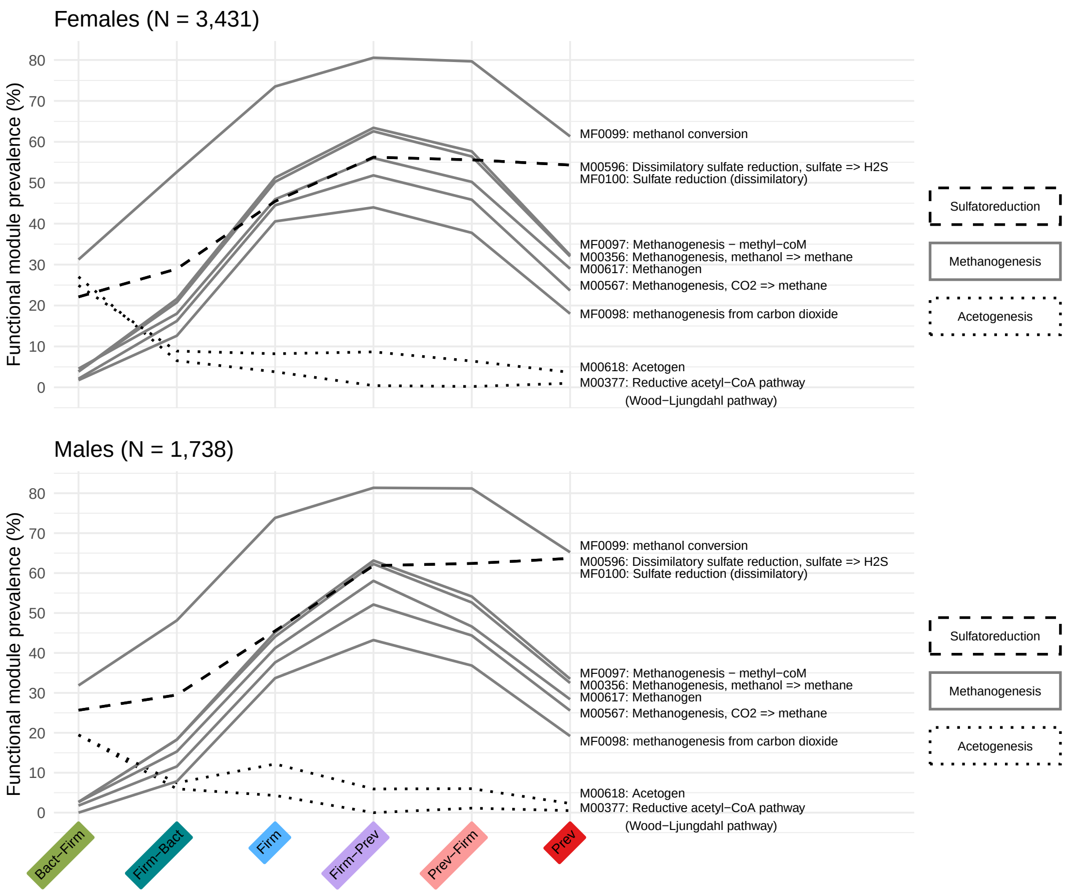


Figure S6: **Prevalence of functional modules linked to $H_2$ across partitions and separated by sex.** The modules' prevalences were evaluated on the LFG data separately on 3,431 females and 1,738 males.

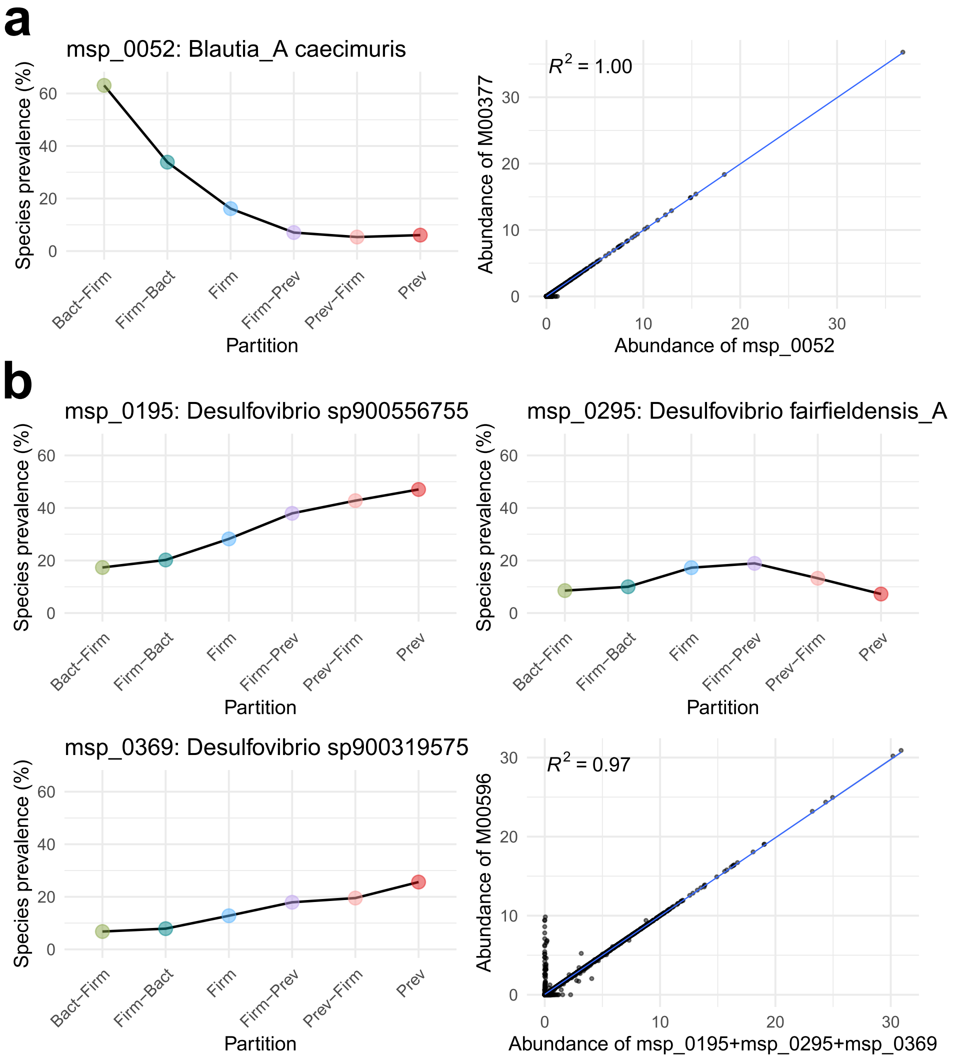


Figure S7: **Genetic evidence supporting hydrogenotrophic sulfate reduction and acetogenesis-associated modules.** a) For acetogenesis (M00377), *Blautia A caecimuris*, the main contributor to this module, also encoded genes associated with $H_2$-dependent $CO_2$ reduction (K22015+K25123+K25124), and its abundance was significantly correlated with that of M00377. b) For dissimilatory sulfate reduction (M00596), three *Desulfovibrio* MSPs carrying the module also encoded $H_2$-oxidizing hydrogenases (K00437+K18008 or K05922+K05927), two of which followed the M00596 prevalence gradient and whose combined abundance was significantly correlated with that of the module. Colors correspond to the partitions color code.

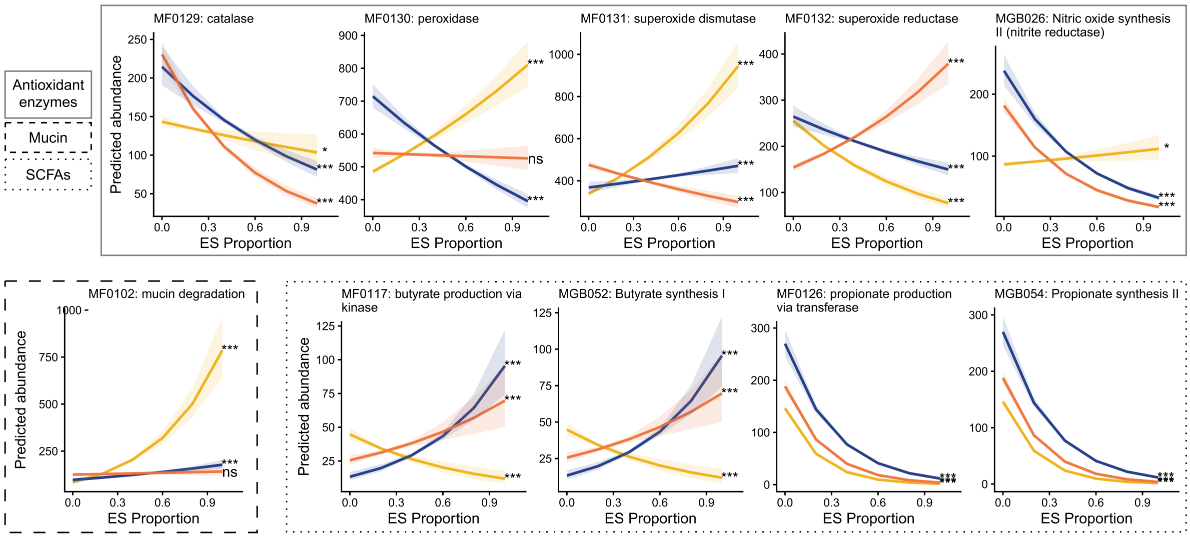


Figure S8: **Prediction of the abundance of modules from Fig. 2b based on the ES proportions.** Modules' abundances were predicted with ES proportions as predictors, using Tweedie generalized linear model (GLM) regressions. The color code is referencing to the three main ES: ES-*Bact* in yellow, ES-*Firm* in blue and ES-*Prev* in orange. The curves correspond to the prediction of the abundance of a module based on 1 ES while the other 2 were considered constant at their sample means. The area around each curve corresponds to a 95 % confidence interval. Asterisks indicate significance of the predictors' adjusted p-values (*p < 0.05, **p < 0.01, ***p < 0.001).

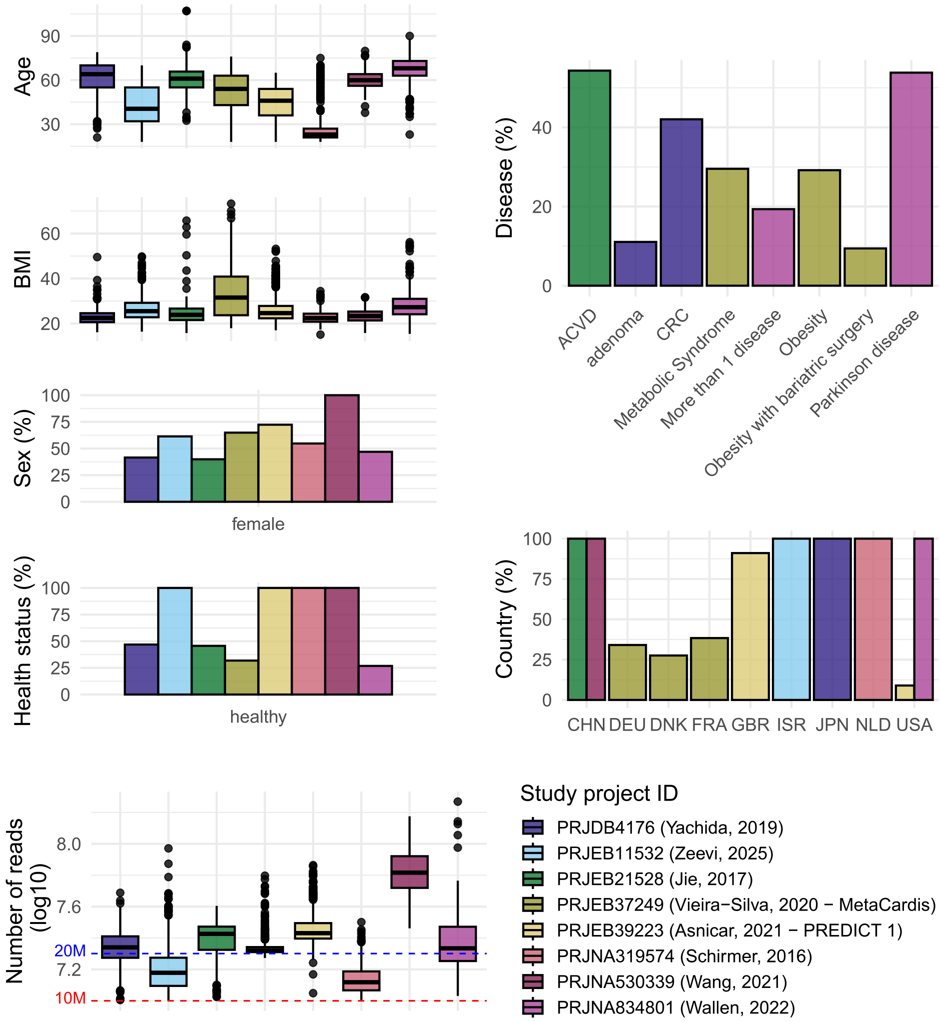


Figure S9: **Meta-cohort characteristics.** Main characteristics and metadata are plotted for each study. For cohorts originally sequenced in paired-end mode, only the forward (R1) reads were retained. The number of reads thus corresponds to single reads. The red dashed line corresponds to 10M single reads, and the blue one to 20M single reads.

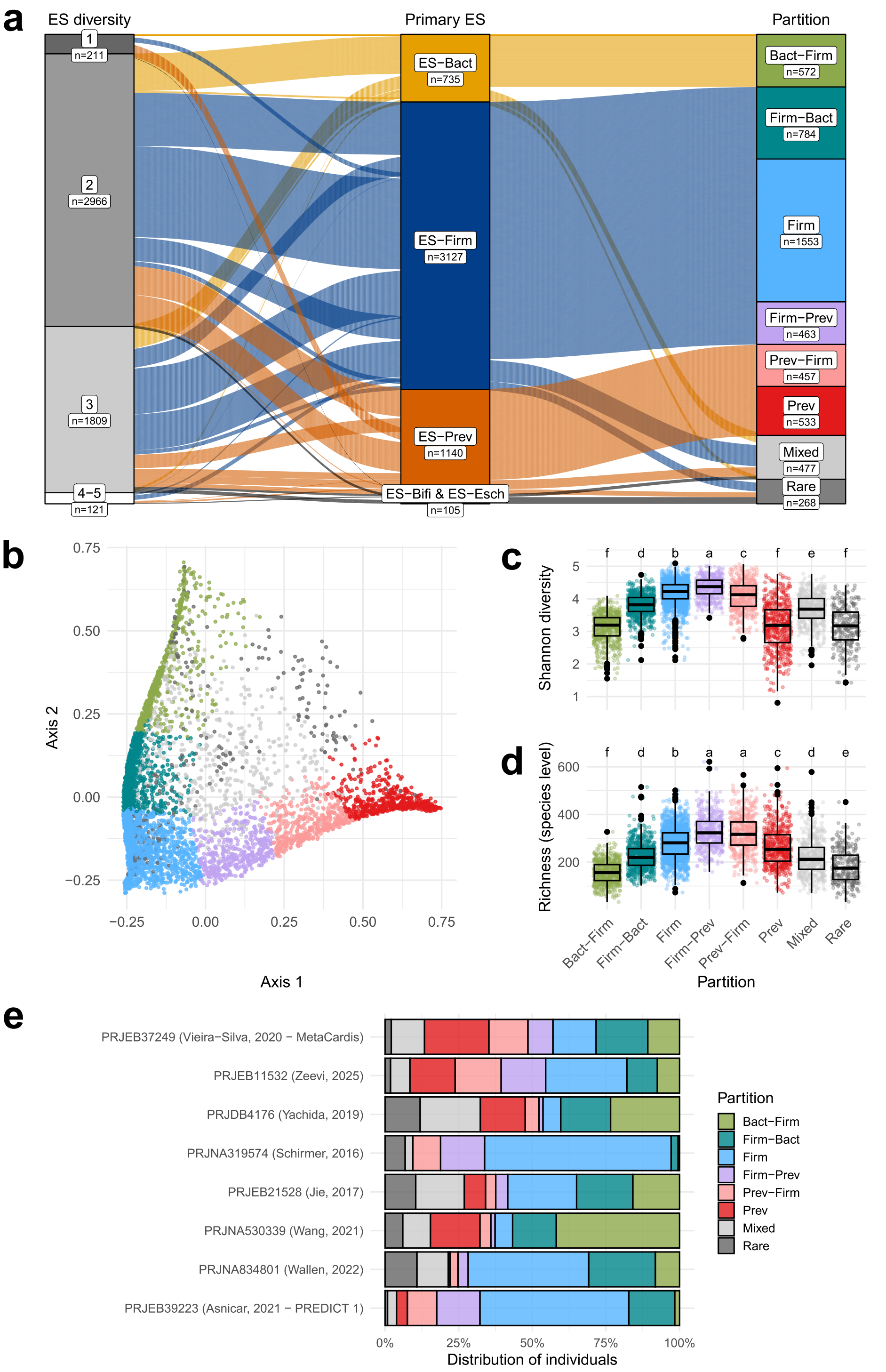


Figure S10: **Enterosignature-based stratification of the international meta-cohort.** a) Individuals distribution across ES and their corresponding partitions. The stratification strategy and the partitioning algorithm are identical to those used for the LFG cohort (see Methods). b) PCoA of Bray-Curtis dissimilarities based on the ES composition of the LFG samples. Each sample is colored according to its partition. c) Distribution of the Shannon diversity of the LFG samples. d) Distribution of the richness of the LFG samples. Letters correspond to the result of Tukey's HSD post-hoc test to assess pairwise significant differences. Different letters indicate that groups are significantly different. e) Distribution of partitions at the study level.

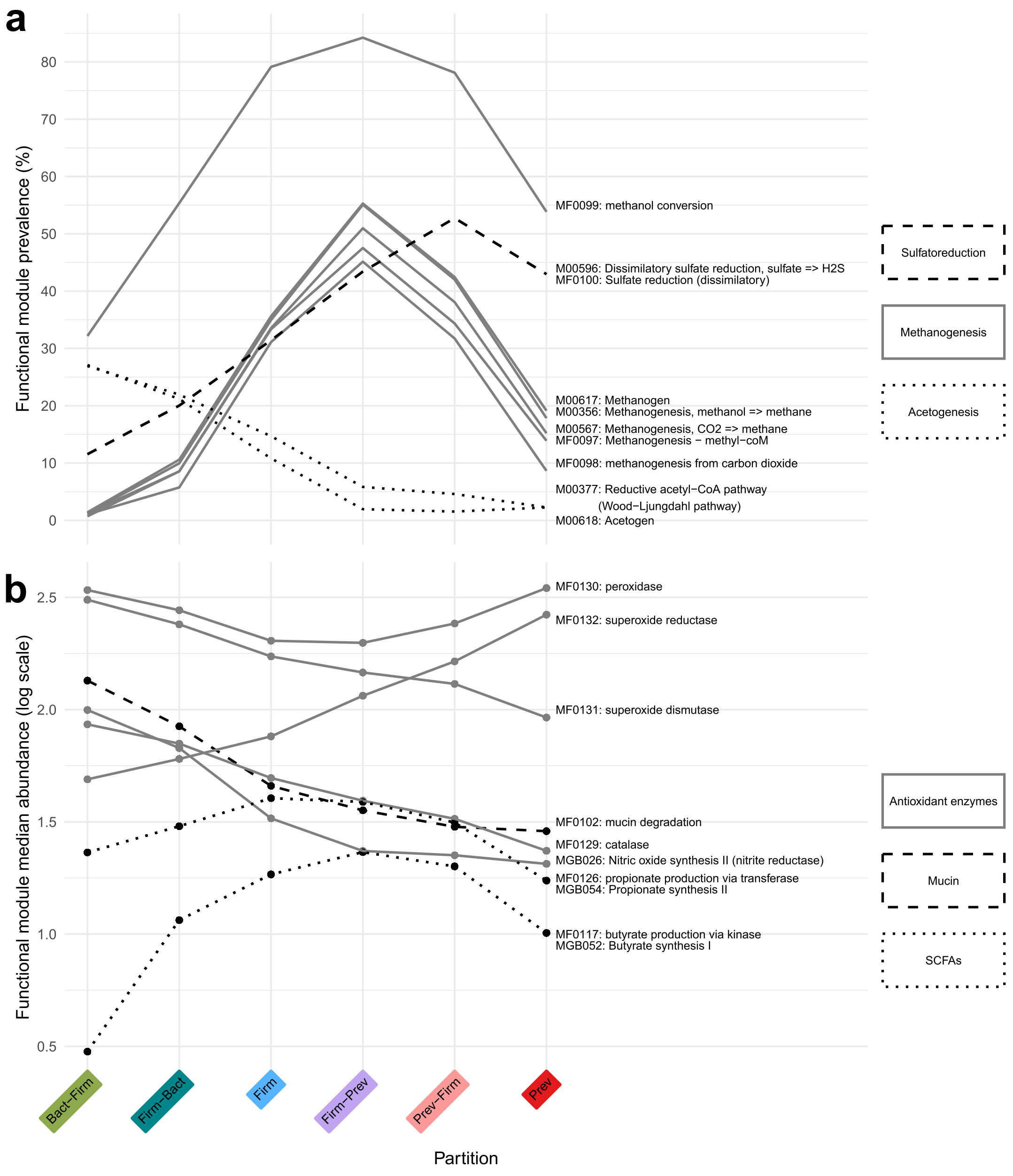


Figure S11: **Trends of functional modules in the meta-cohort.** a) Prevalence of functional modules linked to dihydrogen metabolism across partitions. These modules belong to three categories: sulfatoreduction, acetogenesis, and methanogenesis. b) Abundance of functional modules across partitions. These modules are linked to antioxidant enzymes, mucin degradation and production of butyrate and propionate. The median is plotted.

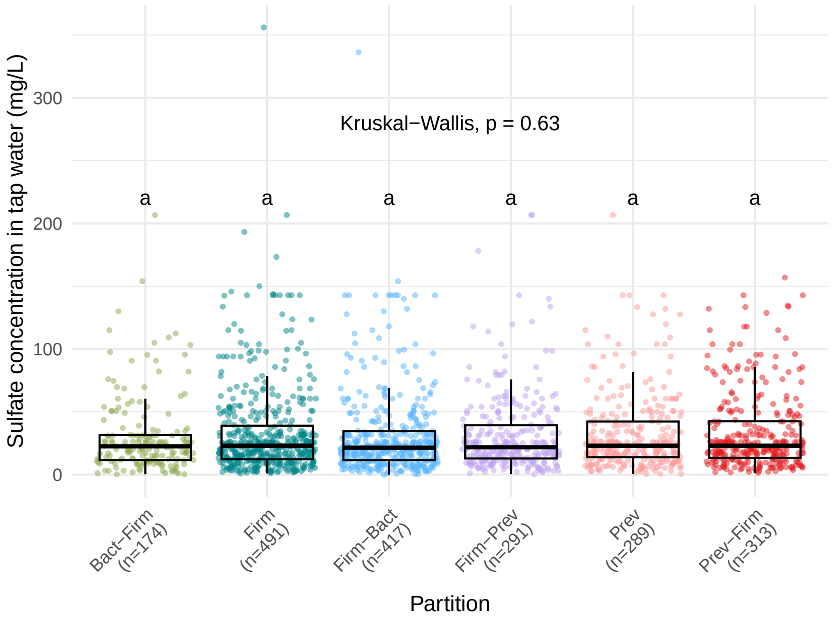


Figure S12: **Sulfate exposition from tap water across partitions.** Each dot corresponds to a LFG participant. The sulfate exposure data extraction is described in Methods.

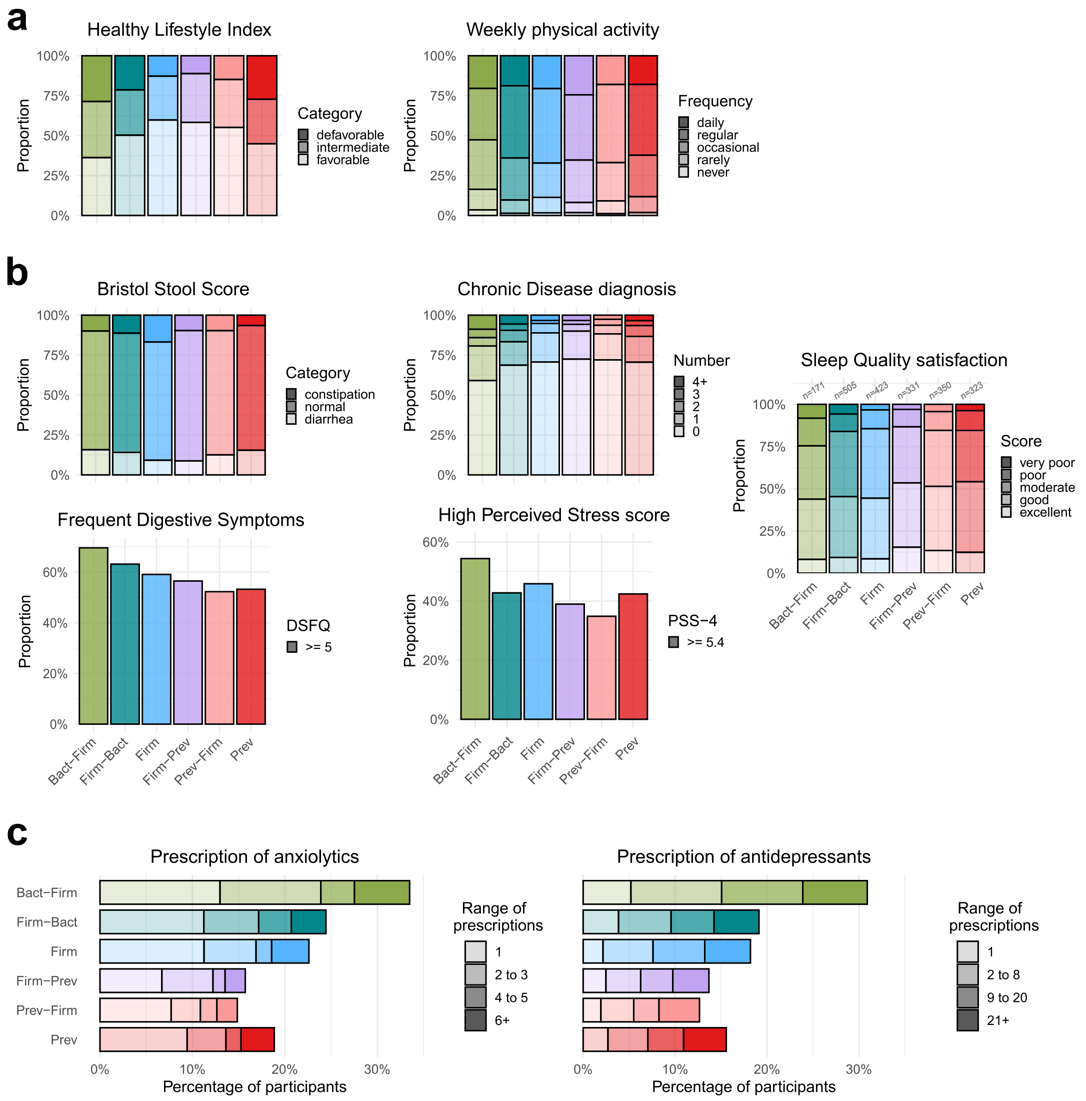


Figure S13: **Health related outcomes across partitions.** a) Distribution of individuals across partitions for lifestyle-related outcomes. b) Distribution of individuals across partitions for health-related outcomes. The frequency of digestive symptoms was considered increased when DSFQ $\geq 5$ [1], and perceived stress elevated when PSS-4 $> 5.4$ [8]. Bristol Stool Scale scores were categorized according to Blake *et al.* [2]. c) Anxiolytic and antidepressants prescriptions across partitions during the two years preceding participation. Colors correspond to the partitions color code, transparency indicates the category level.

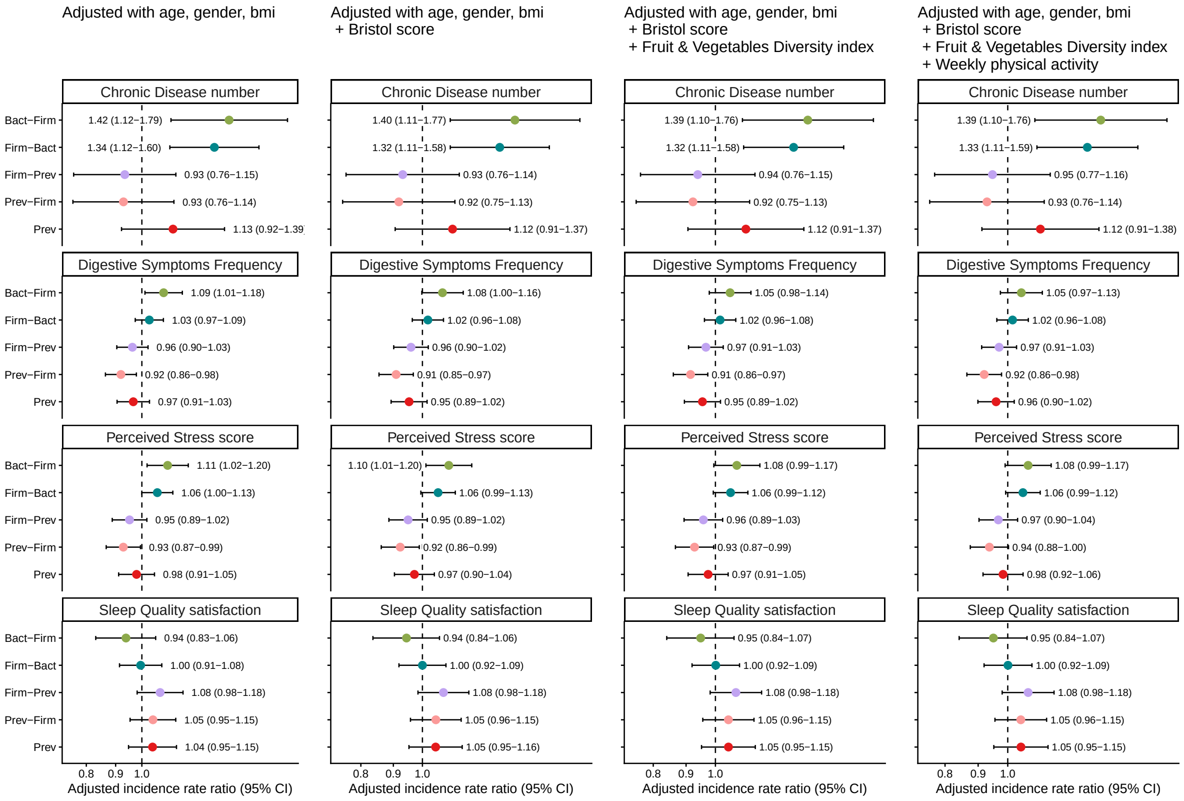


Figure S14: **Associations between partitions and health-related outcomes across sequentially adjusted models.** Forest plots show adjusted incidence rate ratios (IRRs) and 95% confidence intervals. The reference partition was *Firm*. The dashed vertical line indicates an IRR of 1. Colors correspond to the partitions color code.

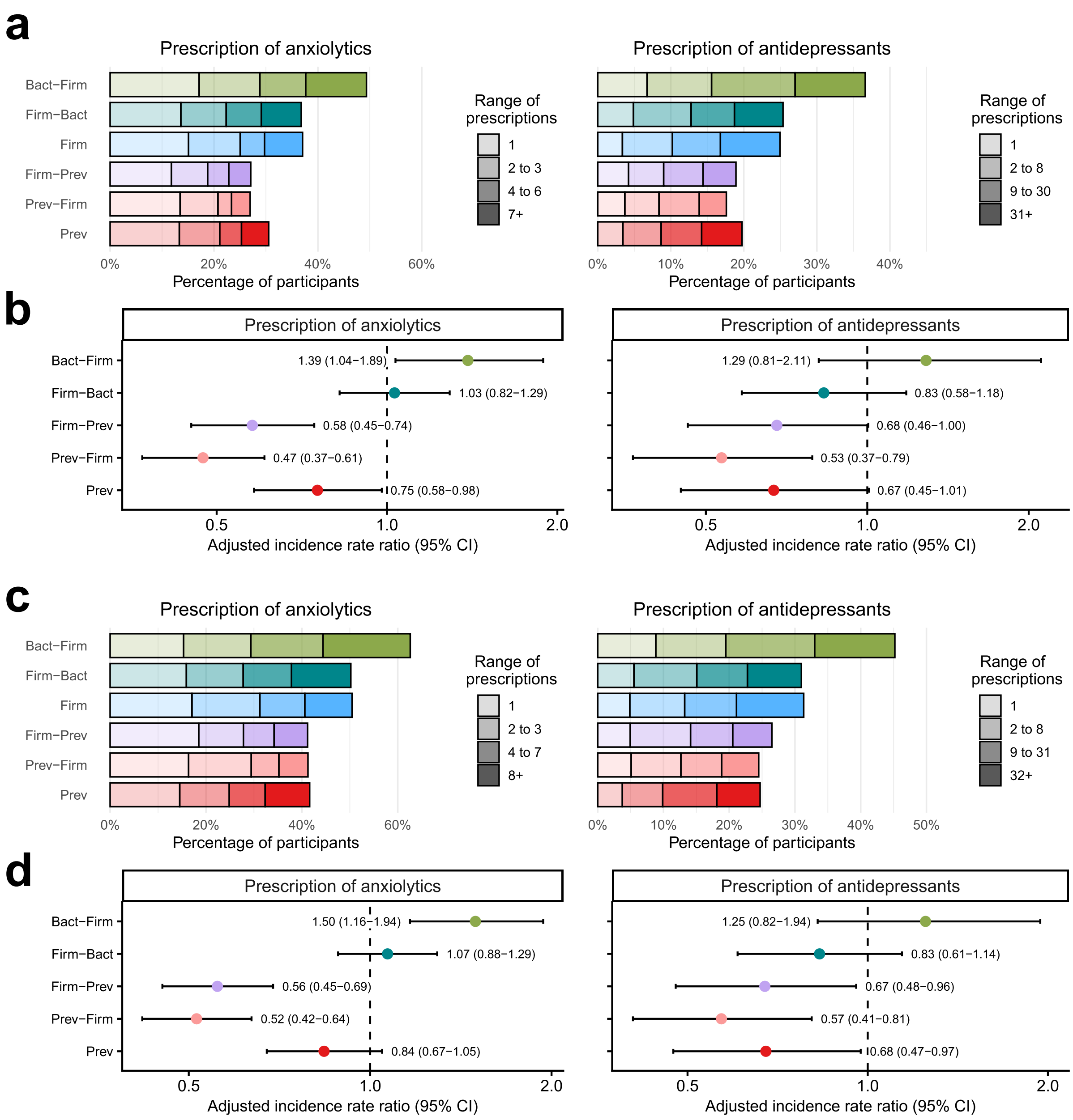


Figure S15: **Psychotropic drug prescription.** a) Anxiolytic and antidepressants prescriptions during the five years preceding participation. b) Associations between partitions and prescriptions for the 5-year window, adjusted for age, sex, and BMI. c) Anxiolytic and antidepressants prescriptions during the ten years preceding participation. d) Associations between partitions and prescriptions for the 10-year window, adjusted for age, sex, and BMI. Forest plots show adjusted incidence rate ratios (IRRs) and 95% confidence intervals; the reference partition was *Firm*. The dashed vertical line indicates an IRR of 1. Colors correspond to the partitions color code, transparency indicates the category level.

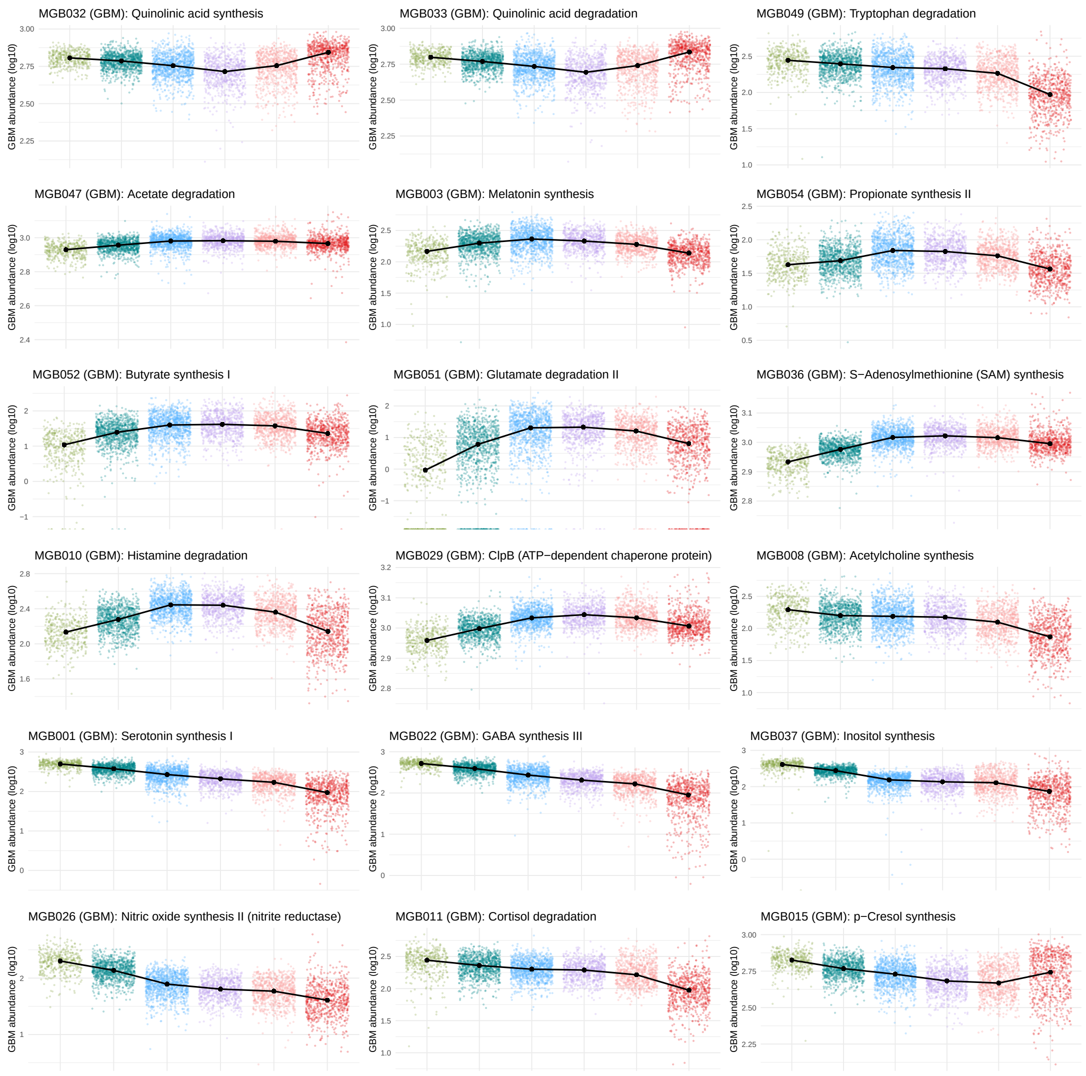


Figure S16: **Gut-Brain modules (GBM) abundance across partitions.** Each dot corresponds to a LFG participant. The curves passes through black dots corresponding to the median for each partition. Plots were reorganized according to the curve trends. The color code is the same as the one previously used to identify partitions. This plot contains all 18 significant functional modules belonging to the GBM database. The selection criteria is detailed in Methods.
